# Translating clinical gene sequencing into a foundational representation of tumor subtype

**DOI:** 10.1101/2025.09.08.674723

**Authors:** JungHo Kong, Ingoo Lee, Dean Boecher, Akshat Singhal, Marcus R. Kelly, Jimin Moon, Chang Ho Ahn, Chan-Young Ock, Tannavee Kumar, Timothy Sears, David Laub, Sarah Wright, Patrick Wall, Hannah Carter, Zhen Wang, Trey Ideker

## Abstract

While gene sequencing is routine in cancer care, translating sequences into treatment decisions remains a challenge. Here we introduce MutationProjector, an AI foundation model that transforms tumor mutation profiles into a compact representation of cancer subtype, with broad implications for diagnosis and therapy. MutationProjector is pre-trained by integrating genomic alterations from >30,000 tumors with extensive molecular knowledge, yielding a model that accurately reconstructs held-out genetic profiles (demonstrating strong generalization) and determines subtype representations from altered molecular pathways (enabling model interpretability). We evaluate MutationProjector in independent tasks related to prediction of immunotherapy response, prediction of chemotherapy response, and classification of metastasis, recording leading performance in all areas. Each task identifies key biomarkers of interest, including *KMT2A* and *KRAS*-*STK11* alterations which govern immunotherapy response.

## Introduction

Tumor genetic profiling is a cornerstone of precision oncology (*1*), enabling patient assessment and treatment decisions across a range of cancer types (*2*). For this purpose, DNA sequencing panels — and in particular those that broadly identify alterations in cancer-associated genes — have been widely adopted in the clinic due to their relatively low cost, rapid turnaround, and established relevance to treatment outcomes (*1*, *3*). Some of the most common clinical gene panels include MSK-IMPACT (*4*, *5*) (400+ cancer-associated genes), FoundationOne CDx (*6*) (324 genes), Tempus xT (*7*) (648 genes), Thermo Fisher Oncomine (161 genes) (*8*), Caris Molecular Intelligence (>700 genes) (*9*), and Guardant360 (70 – 500+ genes depending on version), with one or more of these tests deployed by the vast majority of oncologists and cancer centers (*10*). As of 2025, more than 60 different Federal Drug Administration (FDA)-approved therapies employ cancer gene sequencing as a companion diagnostic. For example, mutation in the *EGFR* gene indicates treatment with gefitinib, erlotinib and others; the BRAF V600E protein variant indicates treatment with vemurafenib or sorafenib, and high tumor mutation burden (TMB) can indicate treatment with anti-PD1/PDL1 immunotherapies.

Despite such progress, the information from genetic sequencing that is clinically actionable remains limited to a few well-studied genes/biomarkers within specific tumor types or therapeutic contexts. Consequently, a match is made to an FDA-approved targeted therapy in only about 8% of cases currently (*11*), usually on the basis of alteration in a single gene. While this situation may reflect the incomplete scope of genes covered by current sequencing panels, it clearly also reflects a fundamental lack of knowledge about how gene mutations should be interpreted. Indeed, the average tumor has approximately 11 distinct genetic alterations identified by clinical sequencing (*12*, *13*), a potentially rich source of molecular information if it could be tapped for therapy selection. One reason that cancer mutations have been difficult to associate with treatment outcomes is that most mutation events are rare (*14*). Another is that individual biomarkers do not function in isolation but act combinatorially to influence a drug response. For instance, while mutations in *ERCC2* (Excision Repair Cross-Complementation Group 2) have been associated with cisplatin sensitivity (*15–17*), drug response may vary depending on other mutations to other DNA repair genes including *BRCA2*, *ATM*, *RB1* and *FANCC* (*18, 19*).

At least two strategies can be applied to address these challenges. First are approaches that integrate cancer mutations with knowledge of molecular networks, since rare and/or co-occurring genetic alteration events are often interconnected within common hallmark pathways (*14*, *20*). Although such analyses are generally based on a single network or pathway resource (*21*), it has been shown repeatedly that integration of multiple networks improves prediction performance in various biological tasks (*21–26*). This improvement can be largely explained by the importance of different types of molecular interactions in tumor pathogenesis and drug response (*27*, *28*), including physical protein-protein binding, protein-DNA transcriptomic regulation and transient signaling modifications including phosphorylation and ubiquitination.

A second approach is to leverage recent developments in artificial intelligence (AI), which present exciting opportunities to harness large-scale cancer genomics data for broader, multi - purpose clinical applications like tumor subtyping and drug response prediction. Foundation models, which are pre-trained on large datasets and then applied to solve diverse new challenges with relatively few samples (*29*, *30*), are especially well positioned to advance precision oncology. Such modeling has been enormously successful in natural language processing (e.g. GPT-5) and computer vision (*29*) and is being actively investigated in life sciences research, including models that analyze DNA (*31*, *32*), RNA (*33–37*), or protein (*38–40*) sequences, or microscopy and radiology images (*41–45*). In oncology in particular, collections of histopathology slides, gene expression profiles, or tumor genomes have been used to train foundation models that were subsequently used to predict somatic mutations (*41*, *46*), cancer type (*41*, *46*), metastatic outcomes (*47*), survival (*47*) or immunotherapy response (*48*, *49*).

Here we present MutationProjector, a machine learning foundation model for decoding cancer genetic alterations. This model is trained from the genetic profiles of more than 30,000 tumor samples gathered across a spectrum of different cancer types (Fig. 1A), integrated with knowledge of cancer molecular networks (Fig. 1B). Once trained, MutationProjector rapidly translates the genome of a tumor into a compact representation of tumor subtype, enabling broad downstream clinical applications.

**Fig. 1.**
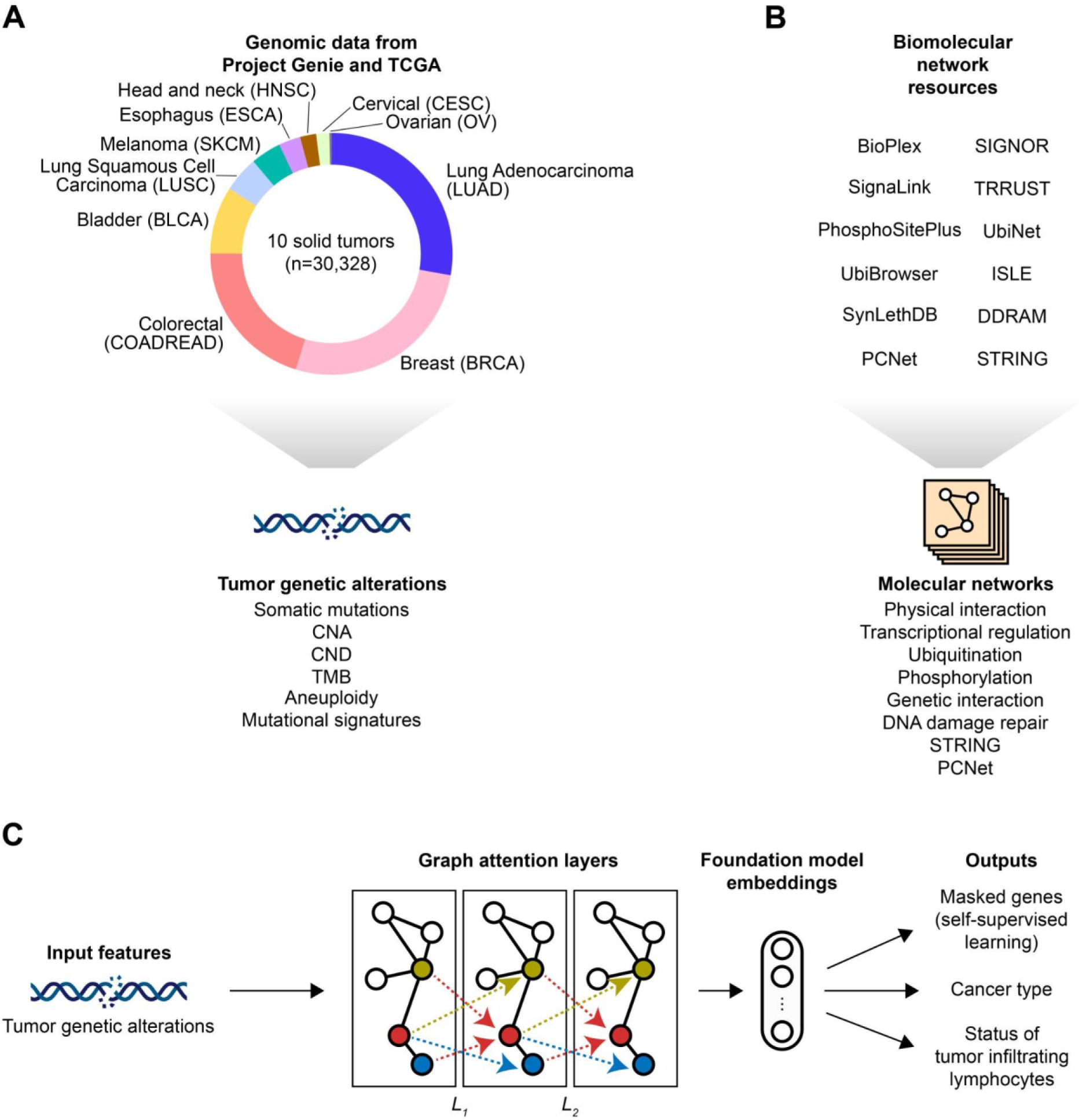
Pre-training on large datasets to enable accurate transfer learning of MutationProjector. (**A**), (**B**) Datasets collected to pre-train MutationProjector, including (**A**) tumor genetic alterations and (**B**) molecular networks. CNA: Copy Number Amplifications, CND: Copy Number Deletions, DDRAM (*67*): DNA Damage Response Assemblies Map, ISLE (*65*): Identification of clinically relevant Synthetic LEthality, PCNet (*23*): Parsimonious Composite Network, SIGNOR (*60*): SIGnaling Network Open Resource, STRING (*68*): Search Tool for Retrieval of Interacting Genes/Proteins, TMB: Tumor Mutation Burden, TRRUST (*61*): Transcriptional Regulatory Relationships Unravelled by Sentence-based Text-mining. (**C**) Overview of MutationProjector pretraining. *L_k_* denotes network-based message-passing at layer *k*.

## Results

### Tumor genetic data and molecular networks

We accessed tumor genetic alterations from Project GENIE (*50*) and The Cancer Genome Atlas (TCGA) (*51*), encompassing 30,328 solid cancers of 10 distinct types (Fig. 1A). From these samples we extracted genetic alterations for 468 genes represented on clinical gene panels, for each gene considering the presence of somatic mutation, copy number amplification or copy number deletion. In addition to these primary genetic data, we included three types of covariates for each tumor: tumor mutation burden (*52*), aneuploidy (*53*) and genome mutational signatures (*54*, *55*).

The average numbers of genes per tumor with somatic mutations, copy number amplifications (CNA) or copy number deletions (CND) were 9.1, 15.9 or 14.0, respectively (sequenced genes only; fig. S1A). Guided by previous reports (*56*, *57*), we verified that certain cancer types display a high rate of somatic mutations and low rate of copy number alterations (e.g. melanoma, bladder cancer) while other cancer types (e.g. ovarian cancer) demonstrate the opposite (fig. S1B). A total of 14 genes had pan-cancer alteration frequencies greater than 10% (considering all types of alterations; fig. S1C).

In parallel to these tumor data, we obtained seven different types of molecular networks, covering general protein-protein physical binding (*58–60*), transcriptional regulation (*59–61*), kinase-substrate phosphorylation (*59*, *60*, *62*), ubiquitin ligase-substrate ubiquitination (*59*, *60*, *63*, *64*), genetic interactions (*65*, *66*), specific protein-protein interactions among DNA damage repair factors (*67*), and global integrated gene-gene networks including STRING (*68*) and PCNet (*23*) (Fig. 1B; Methods). These networks were complementary with respect to pairwise interactions (highest pairwise Jaccard similarity = 0.094; Table S1).

### An integrative foundation model of cancer gene alterations

MutationProjector uses the graph attention network (*69*, *70*) architecture to translate high dimensional data from tumor gene alterations and covariates into a compact low dimensional representation of tumor subtype (Fig. 1C and fig. S2A-C). This representation is created by learning associations among genes, implemented by message-passing within each of the input interaction networks, as well as between genes and covariates (Methods). Associations are learned through attention mechanisms similar to the Transformer model (*71*).

Model optimization was based on three independent pre-training tasks (Fig. 1C). The primary task involved self-supervised learning, in which the model was trained to optimally recover alteration states of masked genes (Fig. 2A). The remaining two tasks were supervised, based on prediction of (i) cancer type and (ii) status of tumor infiltrating lymphocytes (TILs; Methods and fig. S3), respectively. Such a mixed self-supervised/supervised training strategy has been used in many foundation models, including AlphaMissense (*72*) and DNAGPT (*73*).

**Fig. 2.**
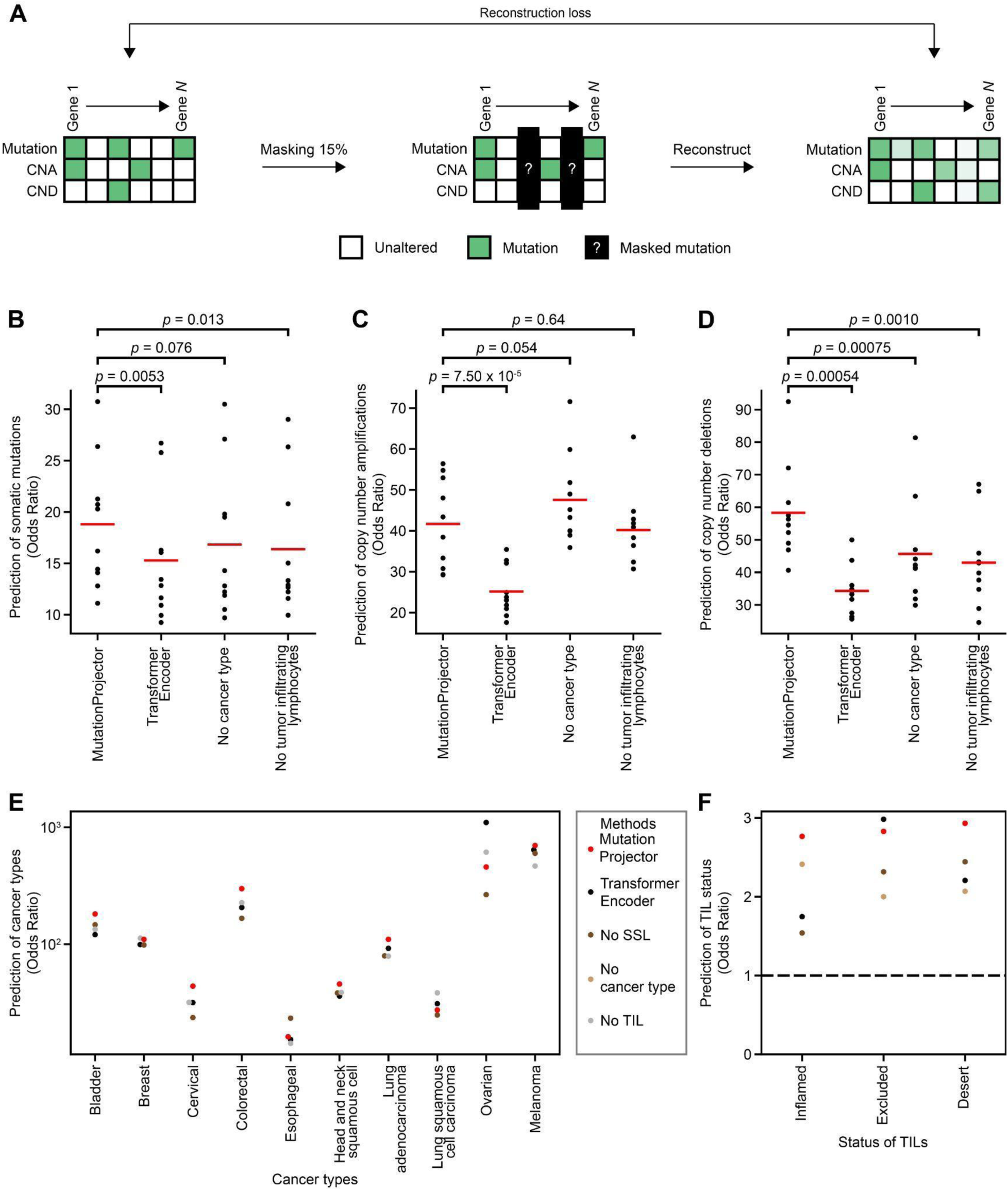
Self-supervised learning and model training performance. (**A**) Schema for self-supervised learning (SSL). At each epoch, 15% of the genes were randomly selected and masked. Reconstruction loss was computed to measure the accurate reconstruction of the masked genes by MutationProjector models. CNA: Copy Number Amplifications, CND: Copy Number Deletions. (**B**)**-**(**D**) Self-supervised learning performance in the held-out (20% of the training dataset) samples. Benchmark models include transformer encoder and models lacking a training task (Methods). 15% of the genes were randomly masked for 10 different iterations to evaluate the prediction of (**B**) somatic mutations, (**C**) CNAs or (**D**) CNDs. Odds ratio, measuring how much actual mutation is in a gene predicted as mutated, was used to estimate the performance of self-supervised learning. Statistical significance was measured using the Mann-Whitney U test. (**E**) Prediction of cancer types in the held-out samples. SSL: Self Supervised Learning, TIL: Tumor Infiltrating Lymphocytes. (**F**) Prediction of immune phenotypes in the held-out samples.

To train and assess the model in each of these tasks, we randomly split the tumor dataset into 80% training (n=24,262) and 20% held-out testing (n=6,066) samples. In the primary self-supervised task, we observed high predictive odds ratios (OR) of 18.8, 41.7 and 58.3 for recovery of held-out somatic mutations (Fig. 2B), CNAs (Fig. 2C) and CNDs (Fig. 2D), respectively. For the remaining two supervised tasks, we found ORs of 16.2 (esophageal cancer) to 698.3 (melanoma) in recovery of each of 10 different cancer types (Fig. 2E); recovery of TIL status was a harder task, with ORs of 2.8, 2.8 or 2.9 for predicting tumors with TIL-inflamed, TIL-excluded or TIL-desert immune landscapes, respectively (Fig. 2F).

We also sought to evaluate the importance of the major aspects of the model architecture and of the training procedure. First, we benchmarked MutationProjector against a naive transformer encoder with a matched number of layers but without any access to network knowledge. MutationProjector displayed significantly better performance in the self-supervised recovery of gene alteration status (Mann-Whitney U test *P* < 0.05 for somatic mutations, CNA and CND; Fig. 2B-D); overall, it also showed better performance in supervised prediction of cancer types (Fig. 2E) and tumor infiltrating lymphocytes (Fig. 2F). Second, we explored substituting the three-task training procedure with simpler training protocols, in which we removed each of the three tasks from training. Most of these dropout experiments resulted in deterioration of predictive performance (Fig. 2B-F), supporting the use of a multi-task training procedure.

### Mutation projections capture complex genetics and subtypes

We next sought to examine and interpret the compact tumor embedding (Fig. 1C) which MutationProjector had learned from the cancer genomic data. Examination of the top two embedding coordinates (UMAP 1 and 2, Fig. 3A) indicated that tumors were approximately stratified by tissue type, with substantial admixture of certain types. For example, tumors from lung squamous cell carcinoma, head-neck squamous cell carcinoma or esophageal cancer were highly similar to one another (Fig. 3B) as part of a larger squamous cluster of tumors (Fig. 3C). Beyond tissue type, the MutationProjector representation was reflective of alteration status of key cancer drivers, such as *APC* mutation in colorectal cancer (*74*), *ARID1A* mutation in a distinct bladder cancer subtype (*75*), *GATA3* mutation in a distinct breast cancer subtype (*76*) or *NFE2L2* mutation in a distinct set of squamous cell carcinomas (*77*) (Fig. 3D). Also discernable in the embedding were distinct clusters of tumors characterized by co-alterations of certain gene pairs involved in genetic or functional interactions, including (i) *FGFR3* mutation and *CDKN2A* deletion (involved in the proliferation-versus-apoptosis axis); (ii) *TP53* and *RB1* mutations (involved in cell-cycle regulation) (*75*); or (iii) *STK11* and *KEAP1* mutations (involved in cellular metabolic and oxidative stress) (*77*) (Fig. 3E). Further inspection of the high attention given by the model to either *FGFR3*-*CDKN2A* co-alteration or *TP53*-*RB1* co-alteration in bladder cancer tumors showed that these events were in fact mutually exclusive (Fig. 3F and Methods), suggesting that MutationProjector had captured distinct molecular subtypes.

**Fig. 3.**
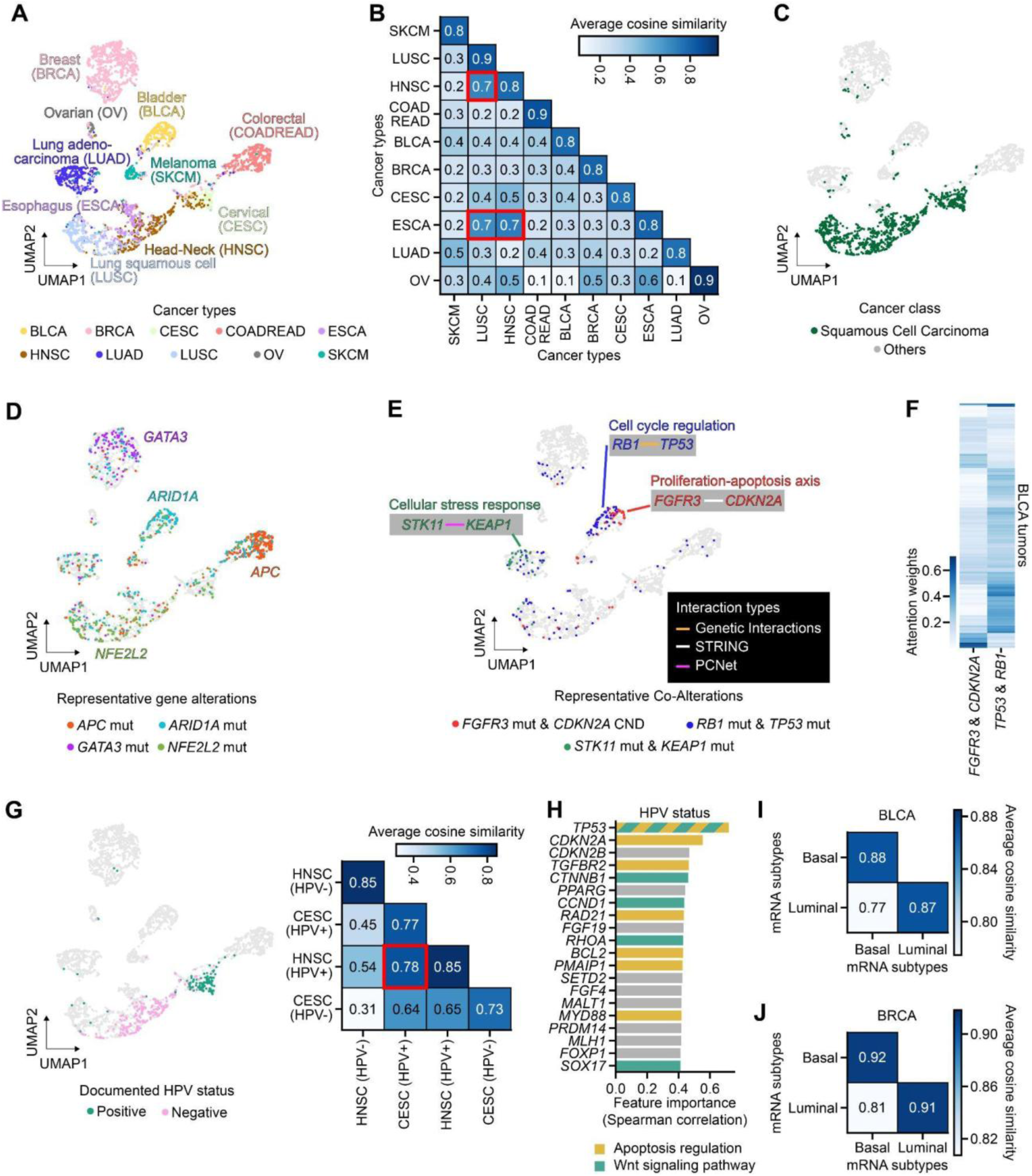
MutationProjector tumor representations capture complex genetics and subtypes. (**A**) Representation of tumors using the final layer embedding of MutationProjector, reduced to two dimensions using the UMAP method (*137*). The points show tumor samples from TCGA (*n*=3,733), colored based on their annotated cancer type. (**B**) Similarity of MutationProjector embeddings across cancer types. Shown for each pair of cancer types is the average of all cosine similarities between the tumors of each pair. Red boxes indicate three tumor type pairs with highest similarity. (**C**) UMAP highlighting squamous cell carcinomas. (**D**) UMAP highlighting tumors with representative genetic alterations. Mut: somatic mutation. (**E**) Representative tumors with co-alteration of gene pairs learned from graph attention. CND: Copy Number Deletion. (**F**) Mutually exclusive attention weights in bladder cancer involving (left column) *FGFR3* and *CDKN2A* or (right column) *TP53* and *RB1*. (**G**) UMAP highlighting the documented Human Papilloma Virus status (HPV, left). Similarities of MutationProjector representations (right). The red box indicates the tumor subtype pair with highest similarity. (**H**) Top 20 features associated with HPV status. Spearman correlation coefficient was used to measure importance. (**I**), (**J**) Similarities of MutationProjector representations between basal and luminal mRNA subtypes in (**I**) bladder and (**J**) breast cancer.

Beyond genetic alteration profiles, we found that the MutationProjector representation had significant correspondence with other types of molecular information not explicitly provided during model training. For example, the embedding identified a cluster of head-and-neck and cervical tumors positive for Human Papilloma Virus (HPV+, Fig. 3G). Although HPV status was not an input to the MutationProjector model, this status had been learned from a coordinated profile of alterations in human genes, including components of the apoptosis regulation or Wnt signaling pathways (e.g. *TP53*, *CDKN2A/B*, *CCND1*; Fig. 3H and Methods) as supported by prior studies (*78–84*). We also found that MutationProjector captured classical tumor subtypes defined formerly by mRNA sequencing, e.g. basal versus luminal transcriptional subtypes in bladder cancer (*75*) (Fig. 3I) or breast cancer (*76*) (Fig. 3J).

### Application in clinical predictive tasks and cohorts

Next, we evaluated the utility of the MutationProjector tumor representation in addressing three independent clinical tasks – prediction of chemotherapy response, prediction of anti-PD1/PDL1-based immunotherapy response, and distinguishing local from metastatic cancers (Fig. 4A). For each of these tasks, we had access to multiple independent cohorts which we each apportioned into training and test sets, comprising *n* = 2,978 patients in all (Table 1). MutationProjector coordinates of each tumor were provided as inputs to a Random Forest classifier for predicting the relevant outcome – sensitivity/resistance to chemotherapy, sensitivity/resistance to immunotherapy, or metastatic/local tumor (Methods).

**Fig. 4.**
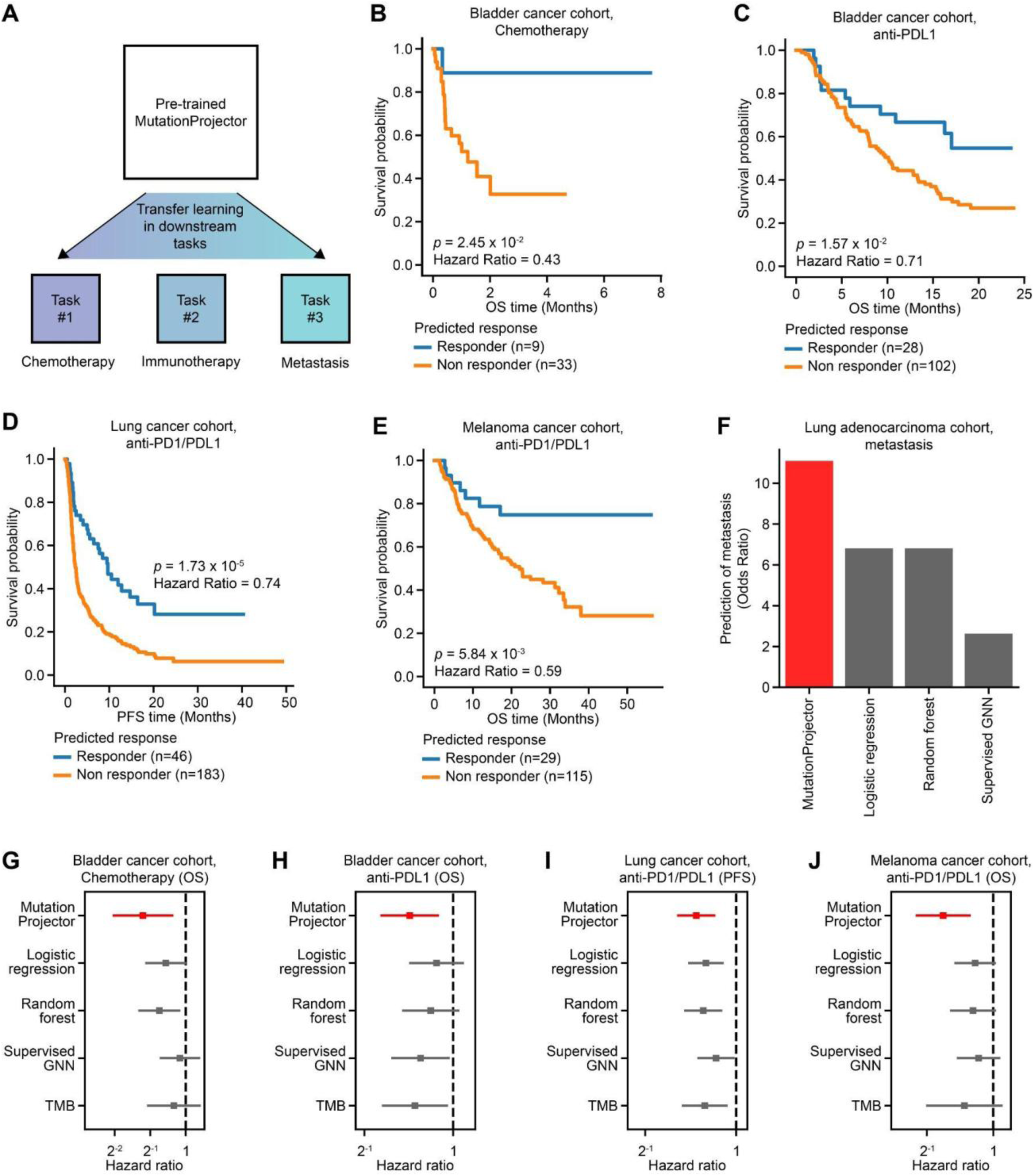
Prediction of drug responses and metastasis in test cohorts. (**A**) Overall scheme for predicting anti-cancer drug treatment responses and metastasis in test cohorts. (**B**)**-**(**E)** Predictive performance of (**B**) chemotherapy response or (**C**)**-**(**E**) anti-PD1/PDL1 response predictions in three independent test cohorts. OS: Overall Survival, PFS: Progression-Free Survival. Statistical significance for survival (either OS or PFS, see x-axis) was measured using the log-rank test. (**F**) Predictive performance of metastasis in an independent test cohort, measured using odds ratios. Other benchmark models include logistic regression, random forest and supervised graph neural network (GNN). (**G**)**-**(**J**) Summary of MutationProjector predictive performance results (red) in comparison to supervised learning approaches (dark grey) in (**G**) chemotherapy and (**H**)**-**(**J**) the three immunotherapy test cohorts. Hazard ratio and its corresponding 95% confidence interval are shown. Dashed vertical line indicates hazard ratio of 1. TMB: Tumor Mutation Burden.

**Table 1.** Summary of downstream tasks and corresponding datasets.

**Datasets used for training**
| Task | Tissue | No. patients | Source |
| --- | --- | --- | --- |
| Chemotherapy | Bladder, Breast, Cervical, Esophagogastric, Head-Neck, Lung, Ovarian, Melanoma | 237 | PMID: 24071849 |
| Immunotherapy | Bladder, Head-Neck, Lung, Melanoma | 94 | PMID: 30150660 |
| Metastasis | Lung adenocarcinoma | 1,974 | PMID: 39506116 |

**Datasets used for testing**
| Task | Tissue | No. patients | Source |
| --- | --- | --- | --- |
| Chemotherapy | Bladder | 42 | PMID: 25092538 |
| Immunotherapy | Bladder | 130 | PMID: 29443960 |
| Immunotherapy | Lung | 229 | PMID: 36038778 |
| Immunotherapy | Melanoma | 144 | PMID: 31792460 |
| Metastasis | Lung adenocarcinoma | 128 | PMID: 37084736 |

For chemotherapy prediction, the classifier was trained on a collection of *n* = 237 cisplatin-treated tumors drawn from eight solid tumor types (*51*), then tested on an independent cohort of *n* = 42 bladder tumors (*85*) (Table 1). Tumors with a predicted chemotherapy response displayed a four month survival rate of 88.9%, in comparison to tumors with a predicted non-response which had a four month survival rate of 32.7% (Fig. 4B). For immunotherapy response prediction, the classifier was trained on anti-PD1/PDL1 treated patients (*n* = 94) encompassing three solid tumor types (*86*), then tested on three independent cohorts representing bladder (*n* = 130) (*87*), lung (*n* = 229) (*88*) or melanoma (*n* = 144) (*89*) cancers (Table 1). In each of these contexts, MutationProjector demonstrated significant prediction of progression-free and overall survival (Fig. 4C-E; log-rank test *P* < 0.05). Finally, for prediction of metastasis we trained the classifier on a lung adenocarcinoma cohort (*n* = 1,974) (*90*) then tested the model in an independent lung adenocarcinoma cohort (*n* = 128) (*91*) (Table 1). Here we observed a very high predictive OR of 11.1 (Fig. 4F). For all downstream tasks evaluated in test cohorts, MutationProjector showed best-in-class performance in comparison to classical supervised learning models or established biomarkers (Fig. 4F-J and fig. S5). For example, in prediction of chemotherapy outcomes MutationProjector embedding-based classifier obtained a low hazard ratio of HR=0.43, indicating strong evidence of benefit, in comparison to the best competing model, Random Forests (using the same genetic alteration information as the MutationProjector), which yielded HR=0.60 (Fig. 4G).

### Clinical tasks integrate mutational patterns across diverse pathways and signatures

Finally, we sought to identify particular genetic biomarkers that had been most important for each of the clinical tasks (fig. S4 and Methods). For chemotherapy response prediction (Fig. 5A), a variety of gene alterations had been deemed important including numerous DNA repair genes, several of which were already well-known markers of chemotherapy response, e.g. *BRCA1* mutation (*92*). Other important features for this task were less well known, including expanded components of DNA repair (e.g. *CHEK1* and *MSH6*) as well as Hedgehog signaling (e.g. *PTCH1* and *GLI1*) and receptor tyrosine kinase pathways (e.g. *ERBB1* and *FGFR1*). Notably, the Hedgehog signaling pathway has been linked to the basal subtype in bladder cancer (*93*, *94*), whereas *FGFR* signaling has been linked to the luminal subtype (*75*), supporting the use of these pathways in prediction of chemotherapy response.

**Fig. 5.**
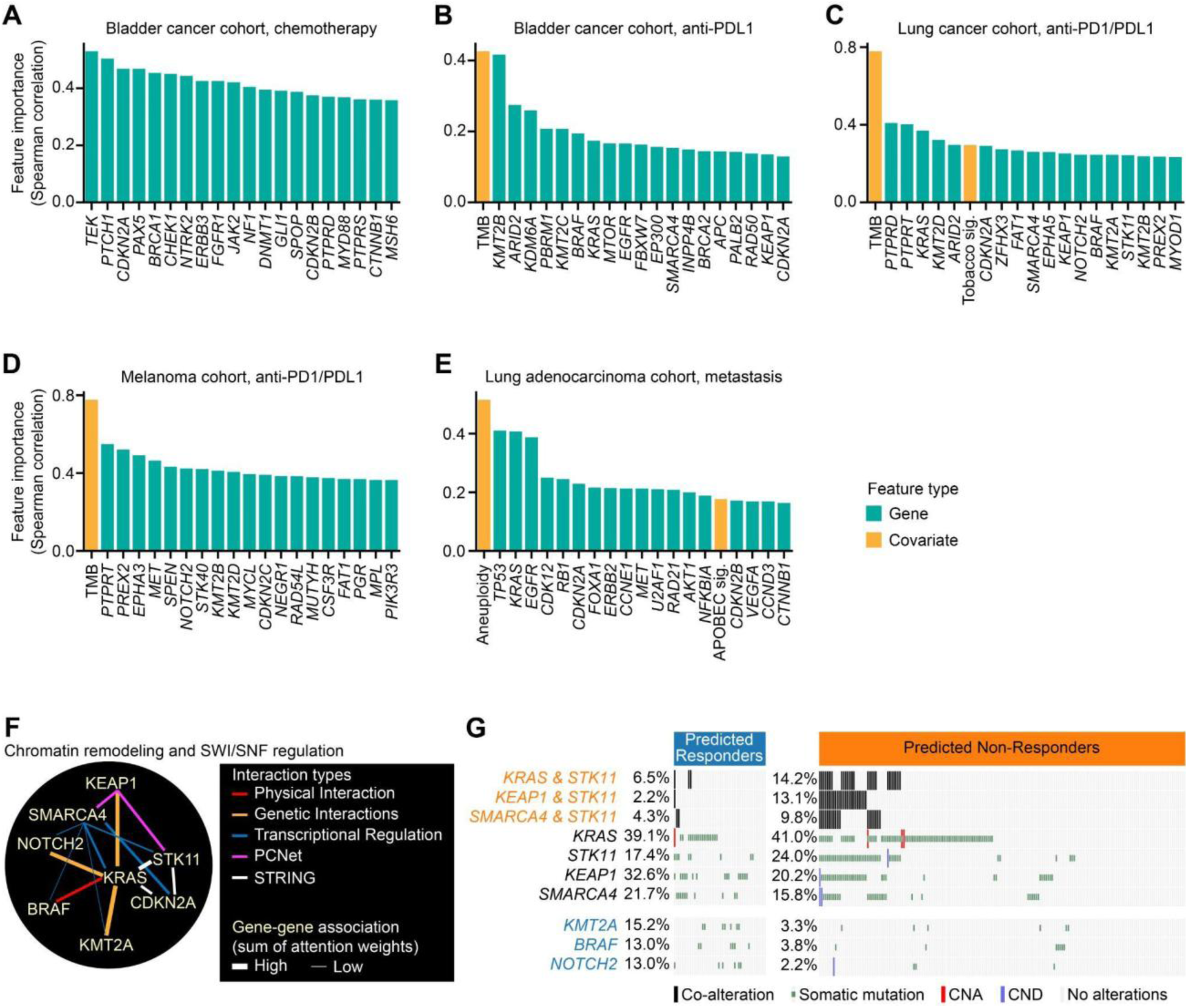
Important features from downstream tasks. (**A**)**-**(**E**) Top 20 important features for (**A**) chemotherapy response prediction in bladder cancer, (**B**) anti-PDL1 response prediction in bladder cancer, (**C**) anti-PD1/PDL1 response prediction in lung cancer, (**D**) anti-PD1/PDL1 response prediction in melanoma and (**E**) metastasis prediction in lung adenocarcinoma. Spearman correlation coefficient was used to measure feature importance. Gene and covariate features were colored teal and orange, respectively. (**F**) Important features in the *KRAS*-centric network (i.e. chromatin remodelling and SWI/SNF regulation-associated genes). Edge color indicates interaction type. Edge thickness indicates the total sum of attention weights on that pairwise gene-gene interaction in the immunotherapy treated lung cohort. (**G**) Alteration patterns of chromatin remodelling and SWI/SNF regulation genes. Genes with significant alterations in the predicted non-responders and responders are shown in orange and blue, respectively (FDR corrected *P* < 0.1). Mutational frequencies in each group are shown as percentages.

Important features for prediction of immunotherapy response included a high tumor mutation burden (TMB) (*95*), which was consistently the top feature in all cohorts including bladder (Fig. 5B), lung (Fig. 5C) and melanoma (Fig. 5D). Unexpected markers for this task included histone methyltransferases (*KMT2A*, *KMT2B*, *KMT2C*), which were strongly predictive of immunotherapy response outcome. Important features for metastasis prediction included genetic alterations in a Wnt signaling pathway gene, *CTNNB1* (*96*), cell-cycle genes, e.g. *CDK12* (*97*) and *CCNE1* (*98*), an angiogenesis-promoting gene, *VEGFA* (*99*), APOBEC mutation signatures (*100*), and aneuploidy (*101*); all of these markers were already well-known to be highly predictive of metastatic tumors (Fig. 5E). Unexpected features of metastasis included components of cell cycle and DNA damage repair (e.g. *CDKN2A*, *CDKN2B* and *RAD21*), a transcription factor (*FOXA1*), and RNA splicing (*U2AF1*).

Among the top 20 important immunotherapy features in lung cancer (Fig. 5C), 7 were alterations in genes functioning in chromatin remodeling and SWI/SNF transcription factors, all of which were involved in physical interaction, transcriptional regulation, genetic or functional interactions with *KRAS*, one of the most highly mutated genes in lung cancer (*102*) (Fig. 5F). Distinct co-alteration patterns in interacting pairs of these genes were observed in predicted non-responders (Fisher’s exact test, FDR corrected *p* < 0.1; Fig. 5G and Table S2), including *KRAS-STK11* (14.2% of predicted non-responders), *KEAP1*-*STK11* (13.1%) or *SMARCA4-STK11* (9.8%). Elsewhere in this network, elevated mutation frequencies were indicative of immunotherapy responders, including *KMT2A* (15.2% of predicted responders), *NOTCH2* (13.0%) and *BRAF* (13.0%) (FDR corrected *p* < 0.1; Table S3).

## Discussion

Here we have explored a strategy for clinical cancer genomics whereby a model of tumor genetic alterations is first ‘pretrained’ from the wealth of tumor DNA sequencing data in the public domain – which are abundant but largely not matched to outcomes – then subsequently tuned for specific clinical tasks using smaller well-annotated cohorts. Following large-scale task-agnostic training on over 30,000 tumor genomes, MutationProjector embeddings were used to accurately predict responses to immunotherapy with further exposure to just 94 samples, achieving leading performance across multiple independent cohorts (Fig. 4G-J). Notably, a single pre-trained tumor model was applied in multiple tasks (1-model / N-tasks), in contrast to the conventional approach of building separate models for each task (N-models / N-tasks). These findings reflect the current movement towards general-purpose AI systems capable of addressing a broad range of challenges (*30*, *103*).

Pretraining the MutationProjector on large genomic data sets allowed the model to capture biologically meaningful cancer subtypes, although the model had not been exclusively trained on these tasks (Fig. 3). First, consistent with a previous report (*104*), MutationProjector generated similar representations of tumor mutations for cancer types that harbor squamous cell carcinomas, displaying high similarities between head-neck cancers, lung squamous cell carcinomas and esophageal cancers (Fig. 3B, C). Moreover, the representations distinguished HPV status in cervical and head-neck cancers (Fig. 3G) with components of apoptosis regulation or Wnt signaling pathway determined as important by MutationProjector (Fig. 3H). In line with our findings, viral oncoproteins (e.g. E6 and E7) encoded by HPV are reported to (i) promote degradation of *TP53* (*105*) to disable apoptosis and checkpoint control, as well as (ii) activate Wnt/beta-catenin signaling pathway (*106*, *107*). In addition to distinguishing HPV status, the representations distinguished basal/luminal mRNA subtypes of bladder and breast cancers (Fig. 3H, I). Notably, basal bladder tumors have been associated with improved survival following neoadjuvant chemotherapy compared to luminal tumors (*108–110*), supporting the use of MutationProjector representations for clinical tasks.

Investigation of MutationProjector’s internal logic revealed a spectrum of individual biomarkers, as well as combinatorial biomarkers within pathways, in which mutational patterns are predictive of immunotherapy response (Fig. 5C and Fig. 5F). Lesser known biomarkers in immunotherapy response included *KMT2A* and *SMARCA4* alterations (Fig. 5G). A potential explanation for these effects is the regulation of the tumor microenvironment through DNA methylation (*111*) or chromatin remodeling (via SWI/SNF complex) (*112*, *113*), with subsequent effects on immunotherapy treatment outcomes. DNA methylation or chromatin remodeling may influence the expression of genes involved in immune cell recruitment or the recognition of tumor cells by the immune system. Indeed, previous studies have highlighted the role of *SMARCA4* in regulating the activity of adaptive immunity-related cytokines such as interferons (*112*, *114*, *115*). Additionally, epigenetic modifications in tumors may alter the activation of genes involved in antigen presentation machinery, including those encoding the Major Histocompatibility Complex (MHC) (*111*, *116*).

Notably, the MutationProjector model was also able to capture combinatorial markers of immunotherapy resistance, such as *STK11-KEAP1* (*117*) or *KRAS-STK11* (*118*) co-alterations, which were used to predict non-responders (Fig. 5G). While mutations in *KRAS*, *STK11* or *KEAP1* were not individually more frequent in non-responders (Table S3), their co-alterations were (Table S2). These results reflect that the embeddings learned by self-supervised approaches can be interpreted to recover complex, multi-gene mutational patterns underlying clinical outcomes.

A further implication is that MutationProjector provides a biologically meaningful embedding of a tumor genome, providing a representation that can be used to identify patients with similar disease. As MutationProjector was trained on clinically actionable genes, patients with similar embeddings should have similar treatment trajectories and may present more effective proxies for projecting likely responses to treatment for a new patient. The pre-trained model could also be used to simulate similar patients with similar tumor characteristics, to serve as digital twins. These simulated patients could then be evaluated for treatment response in the transfer learned models to give a sense of how likely a patient would be to benefit. While a promising direction, it is not yet clear how much data will be required for such efforts.

While 30,000+ genomes representing 10 solid tumor types were considered in our study, numerous additional tumor samples are available for expansion of MutationProjector to tumor types such as pancreatic cancer, prostate cancer or sarcomas (*119–121*). Further cancer genomic resources, such as those provided by The International Cancer Genome Consortium (ICGC) (*122*), may yet further improve predictive performance and biomarker identification. Finally, in addition to genomic data, complementary information from other frequently collected modalities – such as annotations in the electronic health record, images from radiological computed tomography scans and mRNA transcriptomic profiles – may well benefit the pre-training step to increase model performance and interpretability. Other future studies should explore the extent to which the MutationProject concept can be applied to further clinical tasks of interest, including application to liquid biopsies of circulating tumor DNAs for early cancer detection.

## Methods

### Data preparation

Clinical datasets for pre-training were retrieved from Project GENIE (*50*) and TCGA (*51*) databases. To remove potential data leakage, any duplicate samples were removed from analysis. For this study, we focused on 10 solid tumor types including bladder cancer (n=2,734), breast cancer (n=8,163), cervical cancer (n=526), colorectal cancer (n=6,152), esophagus cancer (n=887), head and neck cancer (n=650), lung adenocarcinoma (n=8,445) and squamous cell carcinoma (n=1,454), melanoma (n=1,230) and ovarian cancer (n=87). We used the following OncoTree IDs (*123*) (recorded in the TCGA database) to select patients harboring the 10 tumor types: BLCA, SOC, CEEN, CEMU, CESC, ECAD, ESCA, ESCC, HNSC, LUSC, LUAD, BRCA, BRCNOS, IDC, ILC, IMMC, MBC, SKCM, COAD, MACR, READ. TILs were computed from hematoxylin and eosin stained images in the TCGA dataset using a previously developed ResNet-based deep learning model to predict the status of immune infiltrating lymphocytes (fig. S3) (*124*), constituting immune inflamed (n=1,052), immune excluded (n=1,666) or immune desert (n=1,015) phenotypes (Table S4). Except for the evaluation of pre-training performance (Fig. 2), all 30,328 tumors were used to pre-train the MutationProjector for downstream applications.

As done in previous studies (*125*, *126*), somatic mutations were prepared from 468 genes included in a clinical gene sequencing (MSK-IMPACT (*4*, *5*)). A gene was marked as mutated (“1”) in a tumor sample if it had the following mutation types: missense/nonsense mutations, frameshift insertions/deletions, splice site regions, in-frame insertions/deletions. For TMB we used the reported values; if not available, we calculated the TMB using the Maftools R package (*52*). To calculate arm-level aneuploidy score, we used the ASCETS R package (*53*). To compute mutational signatures, we used the results of the MESiCA algorithm (*55*) for patients sequenced using clinical gene sequencing assay, whereas we used SigProfiler (*54*) for patients with whole-exome/-genome sequencing. As done in a previous study (*55*), we defined seven dominant signatures corresponding to: APOBEC (Apolipoprotein B mRNA editing enzyme catalytic subunit), Clock Signature SBS1, Clock Signature SBS5, Mismatch Repair (MMR), DNA Polymerase Epsilon (POLE), Tobacco, and Ultraviolet light (UV). For APOBEC, Tobacco, UV and MMR, a relative contribution of 30% was labeled as dominant signature (“1”). For Homologous Recombination Deficiency (HRD) and POLE signatures, relative contributions of at least 50% and 20% were used, respectively. For Clock SBS1 and Clock SBS5 signatures, samples with no other signature and at least 40% relative contributions were labeled as having a dominant signature.

### Molecular networks

We considered eight molecular networks in total, encompassing five different molecular interaction types: physical interaction, transcriptional regulation, phosphorylation, ubiquitination, genetic interaction, and, as well as three functional networks (DNA damage repair interaction (*67*), PCNet (*23*) v1.3 and STRING (*68*) v12). For physical interactions, we integrated BioPlex (‘shared’ interaction resource) (*58*), SIGNOR (*60*) and SignaLink (*59*). For transcriptional regulation, we integrated TRRUST (*61*) v2, SIGNOR (*60*) and SignaLink (*59*). For phosphorylation, we integrated PhosphoSitePlus (*62*), SIGNOR (*60*) and SignaLink (*59*). For ubiquitination, we integrated UbiNet 2.0 (*63*), UbiBrowser 2.0 (*64*), SIGNOR (*60*) and SignaLink (*59*). For genetic interactions, we integrated ISLE (*65*) and SynLethDB (*66*). For DNA damage repair we used the DDRAM network (*67*). The NDEx (*127*) repository was used to download PCNet, STRING, BioPlex and DDRAM networks. The number of edges for each network were 1,239 for physical interaction, 3,487 for transcriptional regulation, 774 for phosphorylation, 152 for ubiquitination, 1,245 for genetic interactions, 359 for DNA damage repair, 3,454 for STRING and 15,632 for PCNet.

### Model architecture and pre-training

The MutationProjector design loosely follows the Transformer encoder architecture (*71*), with extensions to encode tumor gene alterations and incorporate molecular networks. Specifically, we used the graph attention network (*69*, *70*), (GAT, instead of a fully connected attention layer as per the original Transformer) to encode network-specific attention knowledge across all samples, enabling biologically-informed decisions (fig. S2A). GAT layers were implemented via GATv2Conv from the pytorch-geometric (*128*) python library. To propagate the impact of local gene-level alteration patterns on various molecular interaction types, we used multi-head attention to create eight attention heads per encoder unit, where each attention head utilized one of the eight distinct interaction networks. This configuration was similar to that of a previous study on driver gene prediction (*25*). For genomic covariates (TMB, aneuploidy and mutational signatures), we made connections from these features to all other nodes in the network in all attention heads, similar to how special tokens or nodes are used in BERT (*129*), the Vision Transformer (*130*) and Graphormer (*131*). This configuration enables the model to aggregate and broadcast genomic covariate information globally, which strengthens long-range interactions via attention patterns. We disabled self-loops in the message passing layer (GAT layer) and applied residual connections as in the original Transformer (*71*) and ResNet (*132*) architectures (fig. S2A). We used two encoder units as done similarly in other graph neural network architectures (*25*, *69*).

Similar to previous self-supervised methods (*33*, *34*, *36*, *129*), we conducted masked gene prediction by hiding (e.g. masking) alteration status of certain genes and asking the model to complete the masked gene’s mutation and copy number alteration status (Fig. 2A). Specifically, each gene is represented as two distinct, learnable vectors, a Gene Token embedding that encodes the gene’s identity (e.g., *TP53*, *KRAS*), and a Mutation Embedding that encodes different combinations of somatic mutations and copy number alternations (fig. S2A). We then randomly select 15% of genes and replace their Mutation Embedding with a special, learnable masked token, while the Gene Token was left untouched. This approach ensures the model knows precisely which gene it needs to make a prediction for, but remains unaware of its actual status. We took element-wise sums of the two token embeddings to generate gene embeddings (fig. S2A). For TMB and aneuploidy, in each cohort, we binned the continuous values into 5 bins, as done similarly in scBERT (*34*). We used 10-dimensional vectors for gene and covariate embeddings (*d*_*features*_; Fig. S1A). To improve prediction performance in the downstream tasks, we borrow ideas from multi - task learning and jointly pre-trained our model on additional supervised labels (i.e. cancer type and levels of TILs) as done similarly in AlphaMissense (*72*) and DNAGPT (*73*). Both cancer types and levels of TILs are categorical labels.

The MutationProjector objective function (Loss) aggregates the prediction errors of self-supervised and supervised learning using binary cross entropy (BCE):

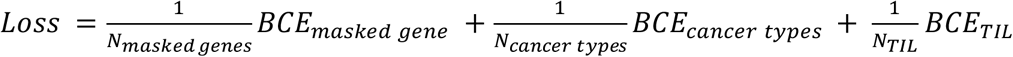

where *N*_*masked genes*_, *N*_*cancer types*_ and *N*_*TIL*_ are equal to the number of genes masked during pre-training, the number of samples with cancer types, and the number of samples with TIL information, respectively. To avoid the tendency to predict abundant classes for the BCE losses, we added weights (inversely proportional to the frequency of a positive label) to label prediction. Hyperparameters such as batch size, number of epochs, learning rate, and dropout rate were set at 64, 100, 0.001, and 0.1, respectively. We optimized the objective function using the AdamW optimizer (*133*), with a weight decay of 0.0001. To determine the importance of (i) networks or (ii) self-supervised/supervised learning, we conducted ablation studies using (i) a fully connected attention layer instead of GAT layer or (ii) removing one supervised learning task or self-supervised learning task during pre-training (Fig. 2B-F). For the fully connected attention layer, we used the same attention layer as in the original Transformer architecture (*71*). All MutationProjector models were implemented in PyTorch and used NVIDIA V100 GPUs.

### Transfer learning and downstream task evaluations

Downstream prediction tasks implemented a transfer learning schema similar to that of scFoundation (*36*), in which the MutationProjector embedding was used as an input for a random forest classifier model (Fig. 4, fig. S2C, 100 trees, max depth 10, class weight = balanced). This random forest that uses MutationProjector embedding was compared to benchmark models including logistic regression, random forest and a supervised graph neural network (similar to MutationProjector but without pretraining), which use gene alterations (mutations and copy number alterations) and genomic covariates as inputs. For the baseline of supervised graph neural network, binary cross entropy loss was used to optimize the model.

To train classifiers on drug response prediction tasks, we used the drug response of each patient to drug treatment as measured according to standard clinical RECIST categories (*134*) (Response Evaluation Criteria in Solid Tumors). We binarized the RECIST categories into responders (Complete Response and Partial Response) and non-responders (Stable Disease and Progressive Disease). For the metastasis prediction task, we used tumor sequencing data from primary tumor biopsies and labeled each patient as metastatic if the pathologic tumor stage was IV, and as non-metastatic if the stage was I (*135*). To allow the MutationProjector to distinguish distal metastasis vs non-metastatic tumor, stage II and III tumors were not included in the analysis.

For downstream prediction tasks (Fig. 4A), we used the classifiers trained for each task and evaluated their predictive performance in independent cohorts. For all downstream tasks, we labeled patients with the top 20% predicted sensitivity scores as ‘responder’ or ‘metastatic’ and otherwise as ‘non responder’ or ‘local tumor’ following thresholds used in prior studies (*95*, *125*). To quantify the performance for predicting drug response and metastasis, we used log-rank test statistics and odds ratios, respectively. Additionally, to assess performance of drug response predictions independent of any cutoff threshold, we computed hazard ratios by running univariate Cox proportional hazards regression using the predicted drug response scores as continuous variables (Fig. 4B-E and Fig. 4G-J). For metastasis prediction, we computed the area under the precision-recall curve (AUPRC) to evaluate the performance of the predictions independent of any cutoff threshold (fig. S6).

### Identification of important features

We used attention weights to quantify the importance of each feature on model predictions. To calculate feature importance, as done in a previous study (*34*) we aggregated multi-head attention weight matrices into a single attention weight matrix by taking an element-wise maximum of the attention weights between a pair of features (fig. S4A, B). This max attention weight matrix was used to identify mutually exclusive attention patterns, for example between *TP53-RB1* and *FGFR3-CDKN2A* in bladder cancer (Fig. 3F). Then, we applied a simplified version of the attention-based linear probing approach that was originally used to investigate BERT (*136*). In detail, to determine the importance of feature *i*, we used a ridge linear regression on the input values of *i*, weighted by incoming and outgoing attention weights, in order to predict model outcomes (fig. S4C). The following function as minimized for each feature:

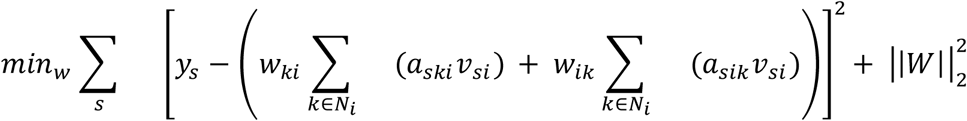

where *y*_*s*_ is the HPV status (Fig. 3H) or the model outcome (Fig. 5) of sample *s*, *k* is the neighbor of feature *i*, *N*_*i*_ is the neighborhood of node *i* in the protein molecular networks, and *v*_*si*_ is the input feature values of *i*. Values *a*_*ski*_ and *a*_*sik*_are incoming and outgoing attention weights between feature *i* and *k*, respectively. Values *w*_*ki*_ and *w*_*ik*_ are learnable parameters of the ridge regression. 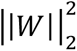 represents L2 regularization. The feature importance was measured by calculating the Spearman correlation between *y* and the predicted outcome of the ridge regression model using feature *i*.

To identify robust important features for drug response and metastasis prediction tasks, we selected features that showed statistically significant importance (Spearman correlation *P* < 0.05) in both training and test cohorts (Table 1). Then in the test cohort, we selected the top 20 important features ranked by the Spearman correlation coefficient (Fig. 5).

## Supplementary Figures

**fig. S1.**
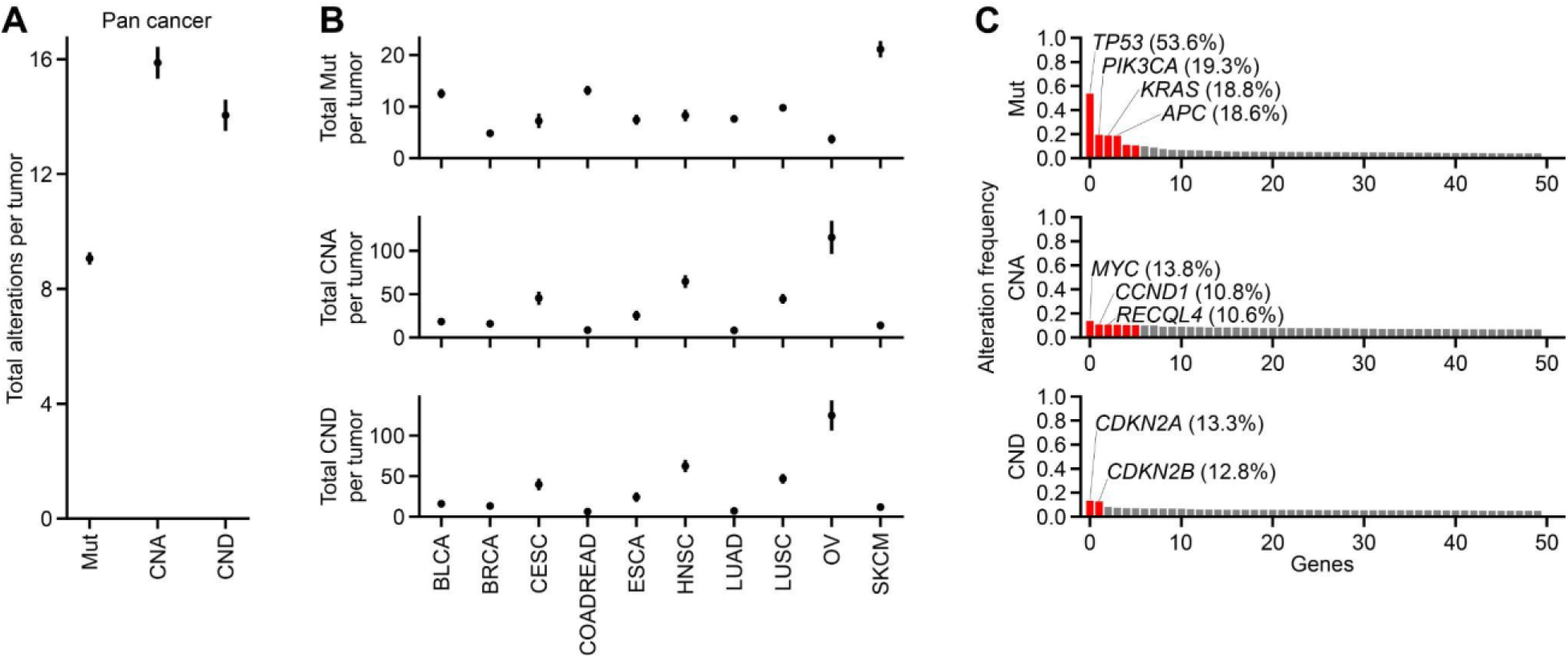
Mutational heterogeneity of the pre-training dataset. **A-C** Total number of gene alterations representing **A** pan cancer alterations and **B** cancer type-specific alterations. Mean and error bars (indicating 95% confidence intervals) are shown. BLCA: Bladder cancer, BRCA: Breast cancer, CESC: Cervical cancer, CNA: Copy Number Amplification, CND: Copy Number Deletion, COADREAD: Colorectal cancer, ESCA: Esophagus cancer, HNSC: Head-Neck Squamous Carcinoma, LUAD: Lung Adenocarcinoma, LUSC: Lung Squamous Cell carcinoma, Mut: somatic mutations, OV: Ovarian cancer, SKCM: Melanoma. **C** Pan cancer gene-level alteration frequencies. Top 50 frequently altered genes per alteration type are shown. Red bars indicate frequencies greater than 10%.

**fig. S2.**
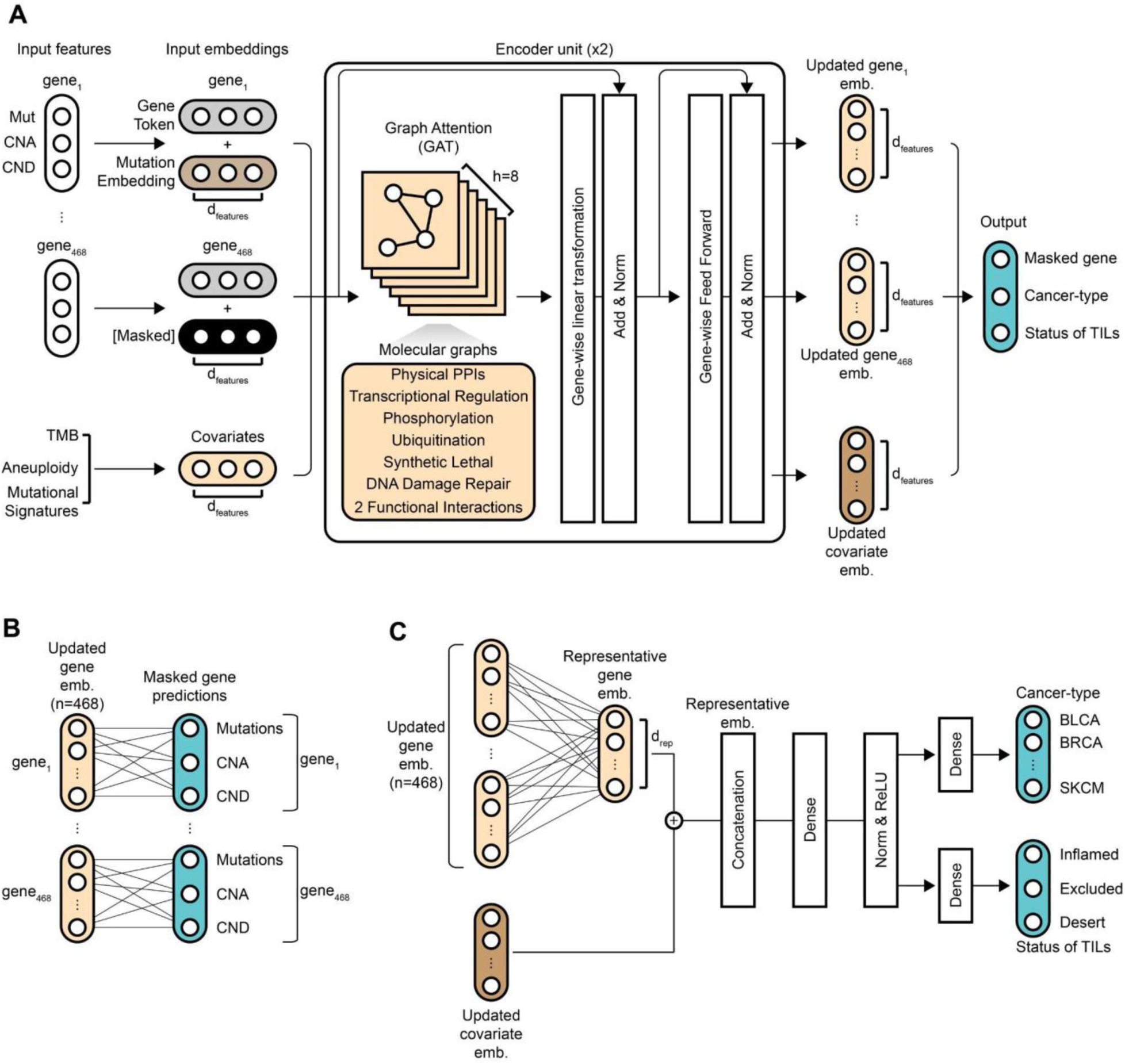
Model architecture. **A** Configuration used during pre-training. Black mutation embedding indicates alteration profiles that are masked. Updated embeddings are generated via network-based message passing using graph attention networks. Size of the gene and covariate embeddings are noted as *d*_*features*_. CNA: Copy Number Amplification, CND: Copy Number Deletion, Emb: Embeddings, Norm: Normalization, TIL: Tumor Infiltrating Lymphocytes, TMB: Tumor Mutational Burden. **B** Self-supervised prediction of masked genes from the updated gene embeddings. **C** Supervised prediction of cancer type and immune infiltration status using the updated gene and covariate embeddings. Size of the representative gene embeddings are noted as *d*_*rep*_. For this study, we set the size of *d*_*rep*_as 90, which matches the total size of the updated covariate embeddings. The dense layer, after the concatenation, has input (i.e. representative embedding) and output sizes of 180 and 10, respectively.

**fig. S3.**
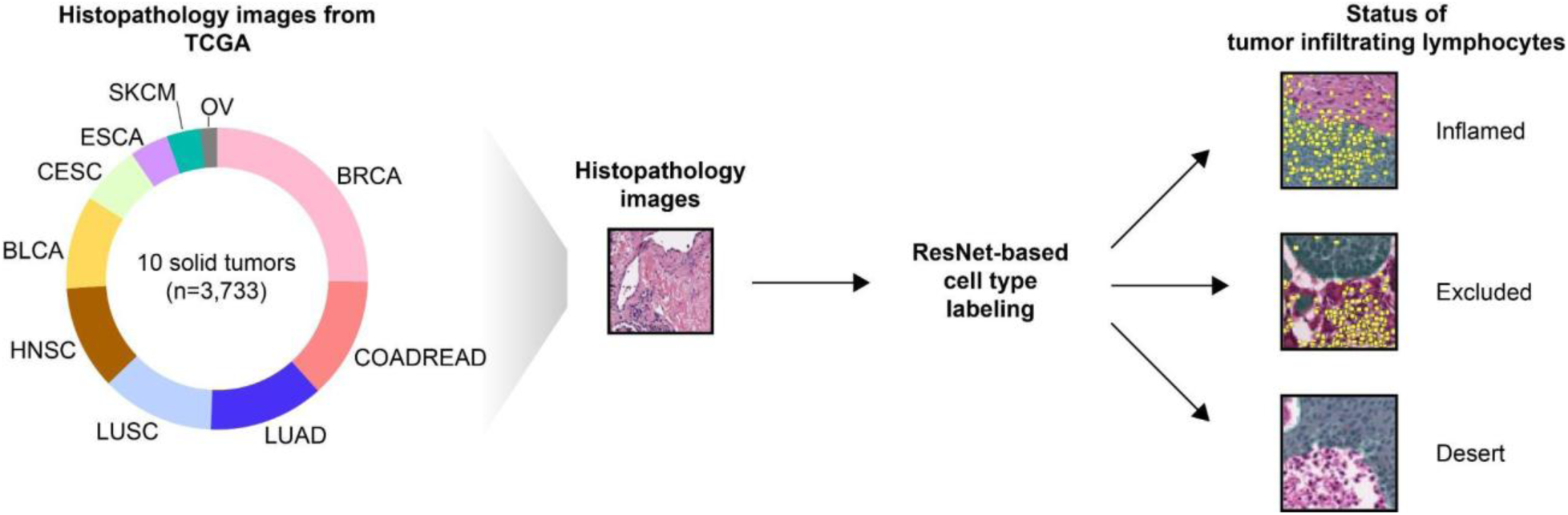
Inferring the status of tumor infiltrating lymphocytes. TIL status is determined from hematoxylin and eosin-stained histopathology image data (left) and classified into one of three labels (far right). BLCA: Bladder cancer, BRCA: Breast cancer, CESC: Cervical cancer, CNA: Copy Number Amplification, CND: Copy Number Deletion, COADREAD: Colorectal cancer, ESCA: Esophagus cancer, HNSC: Head-Neck Squamous Carcinoma, LUAD: Lung Adenocarcinoma, LUSC: Lung Squamous Cell carcinoma, OV: Ovarian cancer, ResNet: Residual neural network, SKCM: Melanoma.

**fig. S4.**
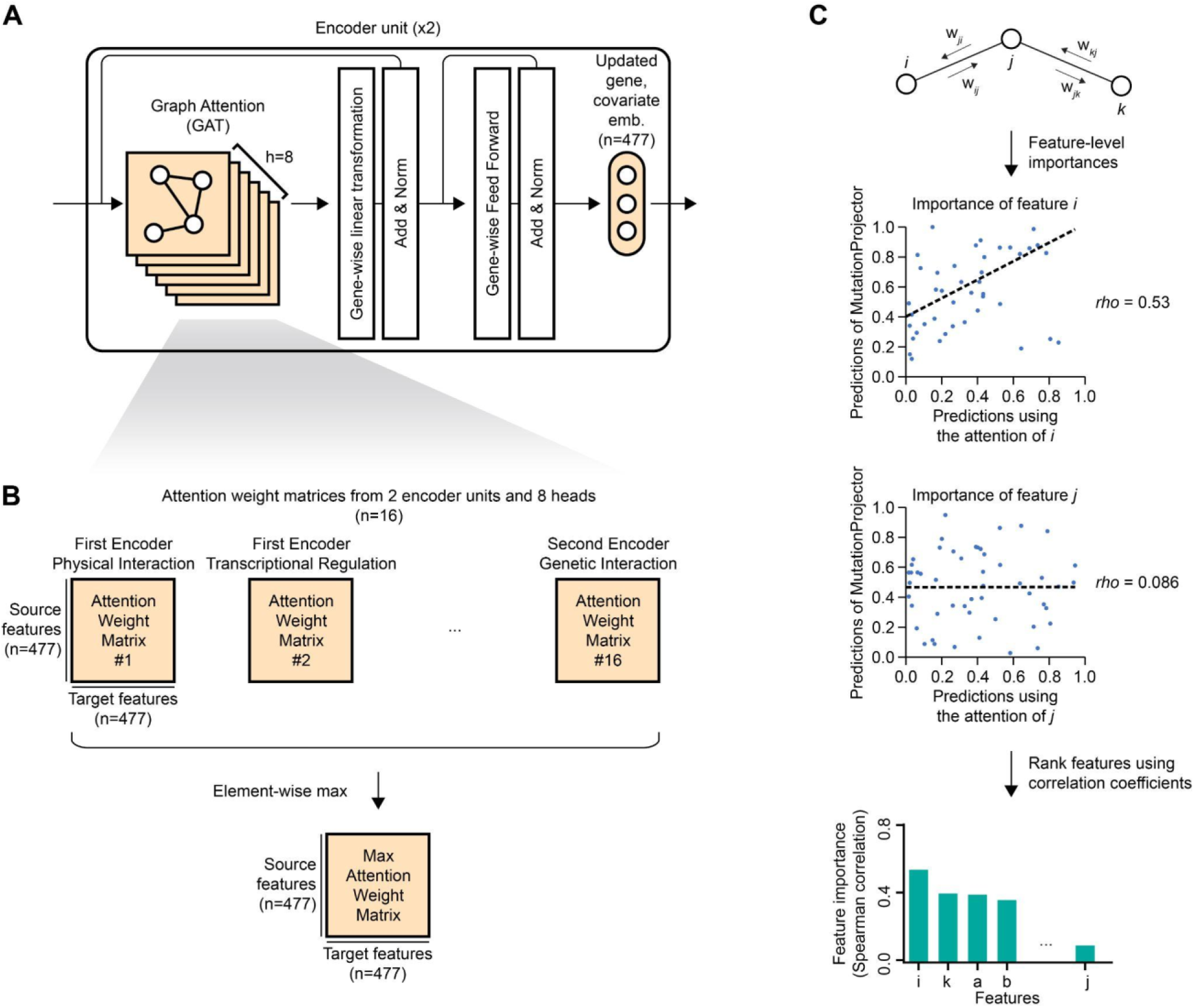
Computing attention weight-based feature importance. **A** Simplified schema of the MutationProjector encoder unit. **B** Summarizing multi-head attention weights in a representative weight matrix. All of the attention weight matrices from 16 different heads were reduced to 1 max attention weight matrix by taking the element-wise max. **C** Computing feature importances using the attention weights from the max attention weight matrix.

**fig. S5.**
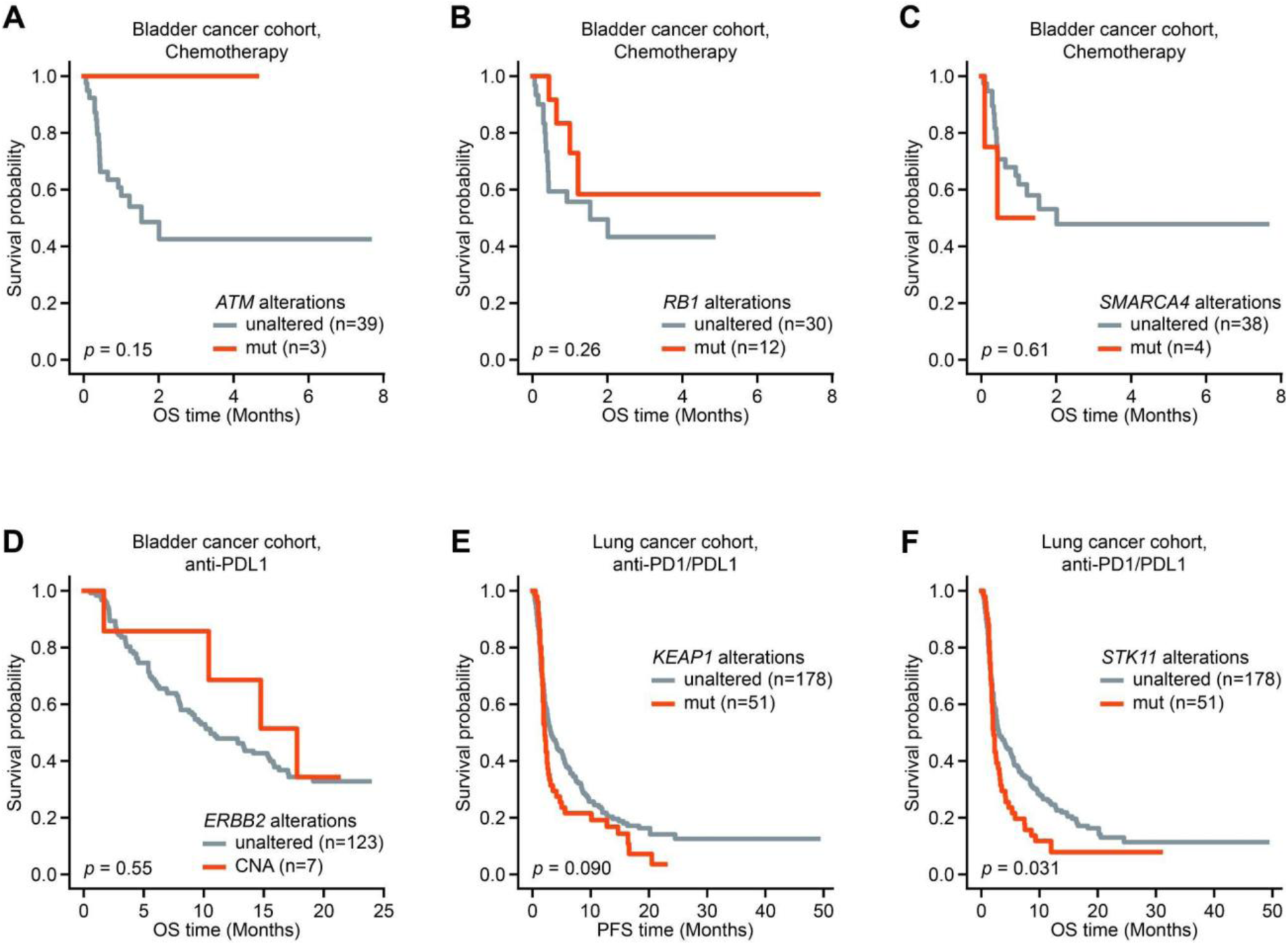
Prediction of drug response using clinical biomarkers. **A-F** Prediction of survival using clinical biomarkers based on **A** *ATM* mutations (*19*), **B** *RB1* mutations (*19*) or **C** *SMARCA4* mutations (*19*) in chemotherapy-treated bladder cancer; **D** *ERBB2* amplifications (*138*, *139*) in immunotherapy-treated bladder cancer; and **E** *KEAP1* mutations (*140*) or **F** *STK11* mutations (*140*) in immunotherapy-treated lung cancer. Statistical significance for a difference in survival between case and control groups was measured using the log-rank test. CNA: Copy Number Amplification.

**fig. S6.**
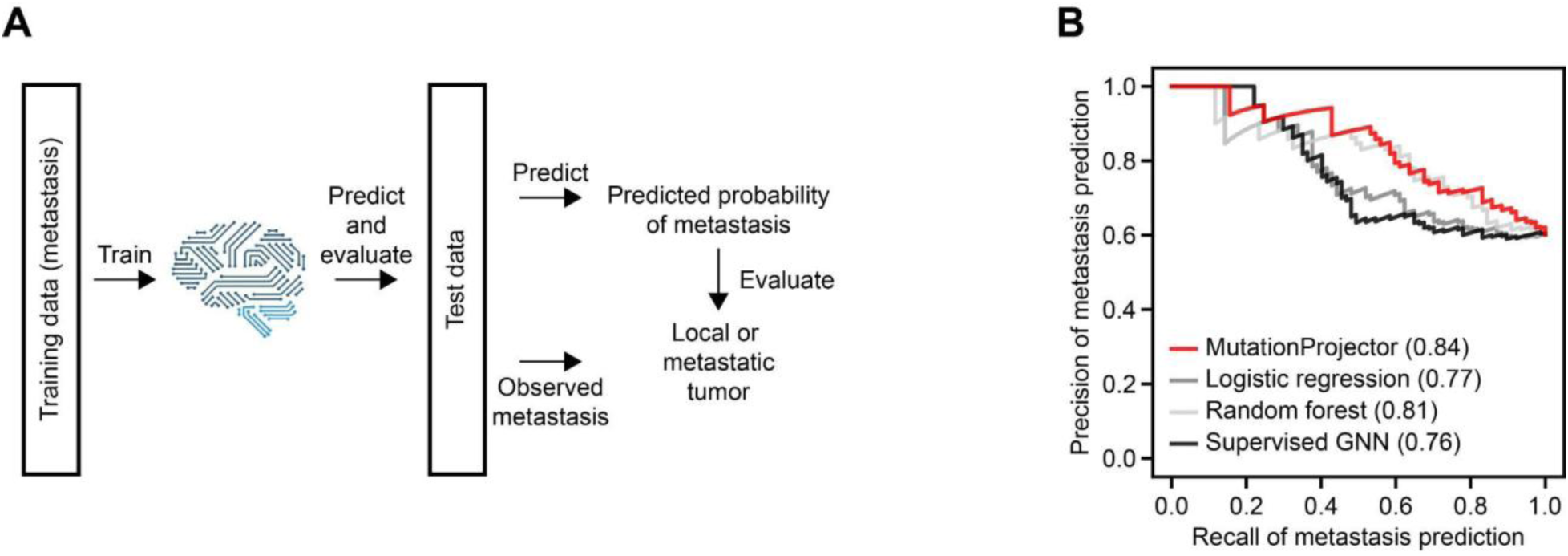
Prediction of metastasis. **A** Overall scheme for predicting metastasis in a test cohort. **B** Predictive performances in the test cohort. The area under the precision-recall curve was used to measure prediction performance. GNN: Graph Neural Network.s

## Acknowledgements

We thank members of the Ideker, Carter and Alexandrov laboratories for helpful discussions. We also acknowledge the data provided by the TCGA Research Network, cBioportal and Project Genie.

## Authors contributions

Conceptualization: JK, IL, MK, TK, DL, SW, HC, ZW, TI

Data curation: JK, DB, AS, TK, TS, PW, HC

Investigation: JK, DB, JM, CA, CO, TK

Visualization: JK, ZW, TI

Funding acquisition: JK, TI

Writing - original draft: JK, IL, AS, MK, HC, ZW, TI

## Data and materials availability

All pharmacogenomics datasets used in this study are publicly accessible through the sources listed (Table 1) (*141–143*). All codes used in this study are available at: https://doi.org/10.5281/zenodo.17073081 and https://github.com/idekerlab/MutationProjector#

## References

1. E. R. Malone, M. Oliva, P. J. B. Sabatini, T. L. Stockley, L. L. Siu, Molecular profiling for precision cancer therapies. Genome Med. 12, 8 (2020).

2. J. K. Sicklick, S. Kato, R. Okamura, M. Schwaederle, M. E. Hahn, C. B. Williams, P. De, A. Krie, D. E. Piccioni, V. A. Miller, J. S. Ross, A. Benson, J. Webster, P. J. Stephens, J. J. Lee, P. T. Fanta, S. M. Lippman, B. Leyland-Jones, R. Kurzrock, Molecular profiling of cancer patients enables personalized combination therapy: the I-PREDICT study. Nat. Med. 25, 744–750 (2019).

3. S. Yip, A. Christofides, S. Banerji, M. R. Downes, I. Izevbaye, B. Lo, A. MacMillan, J. McCuaig, T. Stockley, G. M. Yousef, A. Spatz, A Canadian guideline on the use of next-generation sequencing in oncology. Curr. Oncol. 26, e241–e254 (2019).

4. A. Zehir, R. Benayed, R. H. Shah, A. Syed, S. Middha, H. R. Kim, P. Srinivasan, J. Gao, D. Chakravarty, S. M. Devlin, M. D. Hellmann, D. A. Barron, A. M. Schram, M. Hameed, S. Dogan, D. S. Ross, J. F. Hechtman, D. F. DeLair, J. Yao, D. L. Mandelker, D. T. Cheng, R. Chandramohan, A. S. Mohanty, R. N. Ptashkin, G. Jayakumaran, M. Prasad, M. H. Syed, A. B. Rema, Z. Y. Liu, K. Nafa, L. Borsu, J. Sadowska, J. Casanova, R. Bacares, I. J. Kiecka, A. Razumova, J. B. Son, L. Stewart, T. Baldi, K. A. Mullaney, H. Al-Ahmadie, E. Vakiani, A. A. Abeshouse, A. V. Penson, P. Jonsson, N. Camacho, M. T. Chang, H. H. Won, B. E. Gross, R. Kundra, Z. J. Heins, H.-W. Chen, S. Phillips, H. Zhang, J. Wang, A. Ochoa, J. Wills, M. Eubank, S. B. Thomas, S. M. Gardos, D. N. Reales, J. Galle, R. Durany, R. Cambria, W. Abida, A. Cercek, D. R. Feldman, M. M. Gounder, A. A. Hakimi, J. J. Harding, G. Iyer, Y. Y. Janjigian, E. J. Jordan, C. M. Kelly, M. A. Lowery, L. G. T. Morris, A. M. Omuro, N. Raj, P. Razavi, A. N. Shoushtari, N. Shukla, T. E. Soumerai, A. M. Varghese, R. Yaeger, J. Coleman, B. Bochner, G. J. Riely, L. B. Saltz, H. I. Scher, P. J. Sabbatini, M. E. Robson, D. S. Klimstra, B. S. Taylor, J. Baselga, N. Schultz, D. M. Hyman, M. E. Arcila, D. B. Solit, M. Ladanyi, M. F. Berger, Mutational landscape of metastatic cancer revealed from prospective clinical sequencing of 10,000 patients. Nat. Med. 23, 703–713 (2017).

5. T. Jibiki, H. Nishimura, S. Sengoku, K. Kodama, Regulations, open data and healthcare innovation: A case of MSK-IMPACT and its implications for better cancer care. Cancers (Basel) 13, 3448 (2021).

6. C. A. Milbury, J. Creeden, W.-K. Yip, D. L. Smith, V. Pattani, K. Maxwell, B. Sawchyn, O. Gjoerup, W. Meng, J. Skoletsky, A. D. Concepcion, Y. Tang, X. Bai, N. Dewal, P. Ma, S. T. Bailey, J. Thornton, D. C. Pavlick, G. M. Frampton, D. Lieber, J. White, C. Burns, C. Vietz, Clinical and analytical validation of FoundationOne®CDx, a comprehensive genomic profiling assay for solid tumors. PLoS One 17, e0264138 (2022).

7. N. Beaubier, R. Tell, D. Lau, J. R. Parsons, S. Bush, J. Perera, S. Sorrells, T. Baker, A. Chang, J. Michuda, C. Iguartua, S. MacNeil, K. Shah, P. Ellis, K. Yeatts, B. Mahon, T. Taxter, M. Bontrager, A. Khan, R. Huether, E. Lefkofsky, K. P. White, Clinical validation of the tempus xT next-generation targeted oncology sequencing assay. Oncotarget 10, 2384–2396 (2019).

8. L. K. Vestergaard, D. N. P. Oliveira, T. S. Poulsen, C. K. Høgdall, E. V. Høgdall, Oncomine^TM^ Comprehensive Assay v3 vs. Oncomine^TM^ Comprehensive Assay Plus. Cancers (Basel) 13, 5230 (2021).

9. K. Homicsko, R. Kenneth, A. Voss, O. Michielin, 3356 Triple wild type melanoma profiling in the Caris Molecular IntelligenceTM registry. Eur. J. Cancer 51, S687 (2015).

10. A. N. Freedman, C. N. Klabunde, K. Wiant, L. Enewold, S. W. Gray, K. K. Filipski, N. L. Keating, D. G. B. Leonard, T. Lively, T. S. McNeel, L. Minasian, A. L. Potosky, D. R. Rivera, R. L. Schilsky, D. Schrag, N. I. Simonds, H. M. Sineshaw, J. P. Struewing, G. Willis, J. S. de Moor, Use of next-generation sequencing tests to guide Cancer Treatment: Results from a nationally representative survey of oncologists in the United States. JCO Precis. Oncol. 2, 1–13 (2018).

11. J. Marquart, E. Y. Chen, V. Prasad, Estimation of the percentage of US patients with cancer who benefit from genome-driven oncology. JAMA Oncol. 4, 1093–1098 (2018).

12. M. Takeda, T. Takahama, K. Sakai, S. Shimizu, S. Watanabe, H. Kawakami, K. Tanaka, C. Sato, H. Hayashi, Y. Nonagase, K. Yonesaka, N. Takegawa, T. Okuno, T. Yoshida, S. Fumita, S. Suzuki, K. Haratani, K. Saigoh, A. Ito, T. Mitsudomi, H. Handa, K. Fukuoka, K. Nakagawa, K. Nishio, Clinical application of the FoundationOne CDx assay to therapeutic decision-making for patients with advanced solid tumors. Oncologist 26, e588–e596 (2021).

13. S. Pinet, S. Durand, A. Perani, L. Darnaud, F. Amadjikpe, M. Yon, T. Darbas, A. Vergnenegre, T. Egenod, Y. Simonneau, V. Le Brun-Ly, J. Pestre, L. Venat, F. Thuillier, A. Chaunavel, M. Duchesne, V. Fermeaux, A. Guyot, S. Lacorre, B. Bessette, F. Lalloué, K. Durand, E. Deluche, Clinical management of molecular alterations identified by high throughput sequencing in patients with advanced solid tumors in treatment failure: Real-world data from a French hospital. Front. Oncol. 13, 1104659 (2023).

14. B. Vogelstein, N. Papadopoulos, V. E. Velculescu, S. Zhou, L. A. Diaz, K. W. Kinzler, Cancer genome landscapes. (2013). 10.1126/science.1235122.

15. E. M. Van Allen, K. W. Mouw, P. Kim, G. Iyer, N. Wagle, H. Al-Ahmadie, C. Zhu, I. Ostrovnaya, G. V. Kryukov, K. W. O’Connor, J. Sfakianos, I. Garcia-Grossman, J. Kim, E. A. Guancial, R. Bambury, S. Bahl, N. Gupta, D. Farlow, A. Qu, S. Signoretti, J. A. Barletta, V. Reuter, J. Boehm, M. Lawrence, G. Getz, P. Kantoff, B. H. Bochner, T. K. Choueiri, D. F. Bajorin, D. B. Solit, S. Gabriel, A. D’Andrea, L. A. Garraway, J. E. Rosenberg, Somatic ERCC2 mutations correlate with cisplatin sensitivity in muscle-invasive urothelial carcinoma. Cancer Discov 4, 1140–1153 (2014).

16. A. Sawant, A. Kothandapani, A. Zhitkovich, R. W. Sobol, S. M. Patrick, Role of mismatch repair proteins in the processing of cisplatin interstrand cross-links. DNA Repair (Amst) 35, 126–136 (2015).

17. Q. Li, A. W. Damish, Z. Frazier, D. Liu, E. Reznichenko, A. Kamburov, A. Bell, H. Zhao, E. J. Jordan, S. P. Gao, J. Ma, P. H. Abbosh, J. Bellmunt, E. R. Plimack, J.-B. Lazaro, D. B. Solit, D. Bajorin, J. E. Rosenberg, A. D. D’Andrea, N. Riaz, E. M. Van Allen, G. Iyer, K. W. Mouw, ERCC2 helicase domain mutations confer nucleotide excision repair deficiency and drive cisplatin sensitivity in muscle-invasive bladder cancer. Clin. Cancer Res. 25, 977–988 (2019).

18. A. Taber, E. Christensen, P. Lamy, I. Nordentoft, F. Prip, S. V. Lindskrog, K. Birkenkamp-Demtröder, T. L. H. Okholm, M. Knudsen, J. S. Pedersen, T. Steiniche, M. Agerbæk, J. B. Jensen, L. Dyrskjøt, Molecular correlates of cisplatin-based chemotherapy response in muscle invasive bladder cancer by integrated multi-omics analysis. Nat. Commun. 11, 4858 (2020).

19. E. R. Plimack, R. L. Dunbrack, T. A. Brennan, M. D. Andrake, Y. Zhou, I. G. Serebriiskii, M. Slifker, K. Alpaugh, E. Dulaimi, N. Palma, J. Hoffman-Censits, M. Bilusic, Y.-N. Wong, A. Kutikov, R. Viterbo, R. E. Greenberg, D. Y. T. Chen, C. D. Lallas, E. J. Trabulsi, R. Yelensky, D. J. McConkey, V. A. Miller, E. A. Golemis, E. A. Ross, Defects in DNA repair genes predict response to neoadjuvant cisplatin-based chemotherapy in muscle-invasive bladder cancer. Eur. Urol. 68, 959–967 (2015).

20. D. Hanahan, R. A. Weinberg, Hallmarks of Cancer: The Next Generation - PIIS0092867411001279.pdf. Cell (2011).

21. L. Cowen, T. Ideker, B. J. Raphael, R. Sharan, Network propagation: a universal amplifier of genetic associations. Nat. Rev. Genet. 18, 551–562 (2017).

22. K. Mitra, A.-R. Carvunis, S. K. Ramesh, T. Ideker, Integrative approaches for finding modular structure in biological networks. Nat. Rev. Genet. 14, 719–732 (2013).

23. J. K. Huang, D. E. Carlin, M. K. Yu, W. Zhang, J. F. Kreisberg, P. Tamayo, T. Ideker, Systematic Evaluation of Molecular Networks for Discovery of Disease Genes. Cell Syst 6, 484–495.e5 (2018).

24. S. N. Wright, S. Colton, L. V. Schaffer, R. T. Pillich, C. Churas, D. Pratt, T. Ideker, State of the interactomes: an evaluation of molecular networks for generating biological insights. Mol. Syst. Biol. 21, 1–29 (2025).

25. M. Chatzianastasis, M. Vazirgiannis, Z. Zhang, Explainable Multilayer Graph Neural Network for cancer gene prediction. Bioinformatics 39 (2023).

26. A. Beyer, S. Bandyopadhyay, T. Ideker, Integrating physical and genetic maps: from genomes to interaction networks. 8, 699–710 (2007).

27. P. U. Avila, T. Padvitski, A. C. Leote, H. Chen, J. Saez-Rodriguez, M. Kann, A. Beyer, Gene regulatory networks in disease and ageing. 20, 616–633 (2024).

28. M. Tognetti, A. Gabor, M. Yang, V. Cappelletti, J. Windhager, O. M. Rueda, K. Charmpi, E. Esmaeilishirazifard, A. Bruna, N. de Souza, C. Caldas, A. Beyer, P. Picotti, J. Saez-Rodriguez, B. Bodenmiller, Deciphering the signaling network of breast cancer improves drug sensitivity prediction. Cell Syst. 12, 401–418.e12 (2021).

29. X. Han, Z. Zhang, N. Ding, Y. Gu, X. Liu, Y. Huo, J. Qiu, Y. Yao, A. Zhang, L. Zhang, W. Han, M. Huang, Q. Jin, Y. Lan, Y. Liu, Z. Liu, Z. Lu, X. Qiu, R. Song, J. Tang, J.-R. Wen, J. Yuan, W. X. Zhao, J. Zhu, Pre-trained models: Past, present and future. AI Open 2, 225–250 (2021).

30. R. Bommasani, D. A. Hudson, E. Adeli, R. Altman, S. Arora, S. von Arx, M. S. Bernstein, J. Bohg, A. Bosselut, E. Brunskill, E. Brynjolfsson, S. Buch, D. Card, R. Castellon, N. Chatterji, A. Chen, K. Creel, J. Q. Davis, D. Demszky, C. Donahue, M. Doumbouya, E. Durmus, S. Ermon, J. Etchemendy, K. Ethayarajh, L. Fei-Fei, C. Finn, T. Gale, L. Gillespie, K. Goel, N. Goodman, S. Grossman, N. Guha, T. Hashimoto, P. Henderson, J. Hewitt, D. E. Ho, J. Hong, K. Hsu, J. Huang, T. Icard, S. Jain, D. Jurafsky, P. Kalluri, S. Karamcheti, G. Keeling, F. Khani, O. Khattab, P. W. Koh, M. Krass, R. Krishna, R. Kuditipudi, A. Kumar, F. Ladhak, M. Lee, T. Lee, J. Leskovec, I. Levent, X. L. Li, X. Li, T. Ma, A. Malik, C. D. Manning, S. Mirchandani, E. Mitchell, Z. Munyikwa, S. Nair, A. Narayan, D. Narayanan, B. Newman, A. Nie, J. C. Niebles, H. Nilforoshan, J. Nyarko, G. Ogut, L. Orr, I. Papadimitriou, J. S. Park, C. Piech, E. Portelance, C. Potts, A. Raghunathan, R. Reich, H. Ren, F. Rong, Y. Roohani, C. Ruiz, J. Ryan, C. Ré, D. Sadigh, S. Sagawa, K. Santhanam, A. Shih, K. Srinivasan, A. Tamkin, R. Taori, A. W. Thomas, F. Tramèr, R. E. Wang, W. Wang, B. Wu, J. Wu, Y. Wu, S. M. Xie, M. Yasunaga, J. You, M. Zaharia, M. Zhang, T. Zhang, X. Zhang, Y. Zhang, L. Zheng, K. Zhou, P. Liang, On the opportunities and risks of foundation models, arXiv [cs.LG] (2021). http://arxiv.org/abs/2108.07258.

31. Y. Ji, Z. Zhou, H. Liu, R. V. Davuluri, DNABERT: pre-trained Bidirectional Encoder Representations from Transformers model for DNA-language in genome. Bioinformatics 37, 2112–2120 (2021).

32. G. Benegas, S. S. Batra, Y. S. Song, DNA language models are powerful predictors of genome-wide variant effects. Proc. Natl. Acad. Sci. U. S. A. 120, e2311219120 (2023).

33. C. V. Theodoris, L. Xiao, A. Chopra, M. D. Chaffin, Z. R. Al Sayed, M. C. Hill, H. Mantineo, E. M. Brydon, Z. Zeng, X. S. Liu, P. T. Ellinor, Transfer learning enables predictions in network biology. Nature 618, 616–624 (2023).

34. F. Yang, W. Wang, F. Wang, Y. Fang, D. Tang, J. Huang, H. Lu, J. Yao, scBERT as a large-scale pretrained deep language model for cell type annotation of single-cell RNA-seq data. Nature Machine Intelligence 4, 852–866 (2022).

35. H. Cui, C. Wang, H. Maan, K. Pang, F. Luo, N. Duan, B. Wang, scGPT: toward building a foundation model for single-cell multi-omics using generative AI. Nat. Methods, doi: 10.1038/s41592-024-02201-0 (2024).

36. M. Hao, J. Gong, X. Zeng, C. Liu, Y. Guo, X. Cheng, T. Wang, J. Ma, X. Zhang, L. Song, Large-scale foundation model on single-cell transcriptomics. Nat. Methods, doi: 10.1038/s41592-024-02305-7 (2024).

37. R. Sheinin, R. Sharan, A. Madi, scNET: learning context-specific gene and cell embeddings by integrating single-cell gene expression data with protein-protein interactions. Nat. Methods 22, 708–716 (2025).

38. M. Baek, F. DiMaio, I. Anishchenko, J. Dauparas, S. Ovchinnikov, G. R. Lee, J. Wang, Q. Cong, L. N. Kinch, R. D. Schaeffer, C. Millán, H. Park, C. Adams, C. R. Glassman, A. DeGiovanni, J. H. Pereira, A. V. Rodrigues, A. A. van Dijk, A. C. Ebrecht, D. J. Opperman, T. Sagmeister, C. Buhlheller, T. Pavkov-Keller, M. K. Rathinaswamy, U. Dalwadi, C. K. Yip, J. E. Burke, K. C. Garcia, N. V. Grishin, P. D. Adams, R. J. Read, D. Baker, Accurate prediction of protein structures and interactions using a three-track neural network. Science 373, 871–876 (2021).

39. J. Abramson, J. Adler, J. Dunger, R. Evans, T. Green, A. Pritzel, O. Ronneberger, L. Willmore, A. J. Ballard, J. Bambrick, S. W. Bodenstein, D. A. Evans, C.-C. Hung, M. O’Neill, D. Reiman, K. Tunyasuvunakool, Z. Wu, A. Žemgulytė, E. Arvaniti, C. Beattie, O. Bertolli, A. Bridgland, A. Cherepanov, M. Congreve, A. I. Cowen-Rivers, A. Cowie, M. Figurnov, F. B. Fuchs, H. Gladman, R. Jain, Y. A. Khan, C. M. R. Low, K. Perlin, A. Potapenko, P. Savy, S. Singh, A. Stecula, A. Thillaisundaram, C. Tong, S. Yakneen, E. D. Zhong, M. Zielinski, A. Žídek, V. Bapst, P. Kohli, M. Jaderberg, D. Hassabis, J. M. Jumper, Accurate structure prediction of biomolecular interactions with AlphaFold 3. Nature 630, 493–500 (2024).

40. M. Baek, R. McHugh, I. Anishchenko, H. Jiang, D. Baker, F. DiMaio, Accurate prediction of protein-nucleic acid complexes using RoseTTAFoldNA. Nat. Methods 21, 117–121 (2024).

41. S. J. Wagner, D. Reisenbüchler, N. P. West, J. M. Niehues, J. Zhu, S. Foersch, G. P. Veldhuizen, P. Quirke, H. I. Grabsch, P. A. van den Brandt, G. G. A. Hutchins, S. D. Richman, T. Yuan, R. Langer, J. C. A. Jenniskens, K. Offermans, W. Mueller, R. Gray, S. B. Gruber, J. K. Greenson, G. Rennert, J. D. Bonner, D. Schmolze, J. Jonnagaddala, N. J. Hawkins, R. L. Ward, D. Morton, M. Seymour, L. Magill, M. Nowak, J. Hay, V. H. Koelzer, D. N. Church, TransSCOT consortium, C. Matek, C. Geppert, C. Peng, C. Zhi, X. Ouyang, J. A. James, M. B. Loughrey, M. Salto-Tellez, H. Brenner, M. Hoffmeister, D. Truhn, J. A. Schnabel, M. Boxberg, T. Peng, J. N. Kather, Transformer-based biomarker prediction from colorectal cancer histology: A large-scale multicentric study. Cancer Cell 41, 1650–1661.e4 (2023).

42. C. Ma, W. Tan, R. He, B. Yan, Pretraining a foundation model for generalizable fluorescence microscopy-based image restoration. Nat. Methods, doi: 10.1038/s41592-024-02244-3 (2024).

43. Y. Zhou, M. A. Chia, S. K. Wagner, M. S. Ayhan, D. J. Williamson, R. R. Struyven, T. Liu, M. Xu, M. G. Lozano, P. Woodward-Court, Y. Kihara, UK Biobank Eye & Vision Consortium, A. Altmann, A. Y. Lee, E. J. Topol, A. K. Denniston, D. C. Alexander, P. A. Keane, A foundation model for generalizable disease detection from retinal images. Nature 622, 156–163 (2023).

44. C. Kim, S. U. Gadgil, A. J. DeGrave, J. A. Omiye, Z. R. Cai, R. Daneshjou, S.-I. Lee, Transparent medical image AI via an image–text foundation model grounded in medical literature. Nat. Med. 30, 1154–1165 (2024).

45. J. Ma, Y. He, F. Li, L. Han, C. You, B. Wang, Segment anything in medical images. Nat. Commun. 15, 654 (2024).

46. H. Xu, N. Usuyama, J. Bagga, S. Zhang, R. Rao, T. Naumann, C. Wong, Z. Gero, J. González, Y. Gu, Y. Xu, M. Wei, W. Wang, S. Ma, F. Wei, J. Yang, C. Li, J. Gao, J. Rosemon, T. Bower, S. Lee, R. Weerasinghe, B. J. Wright, A. Robicsek, B. Piening, C. Bifulco, S. Wang, H. Poon, A whole-slide foundation model for digital pathology from real-world data. Nature 630, 181–188 (2024).

47. S. Azizi, L. Culp, J. Freyberg, B. Mustafa, S. Baur, S. Kornblith, T. Chen, N. Tomasev, J. Mitrović, P. Strachan, S. S. Mahdavi, E. Wulczyn, B. Babenko, M. Walker, A. Loh, P.-H. C. Chen, Y. Liu, P. Bavishi, S. M. McKinney, J. Winkens, A. G. Roy, Z. Beaver, F. Ryan, J. Krogue, M. Etemadi, U. Telang, Y. Liu, L. Peng, G. S. Corrado, D. R. Webster, D. Fleet, G. Hinton, N. Houlsby, A. Karthikesalingam, M. Norouzi, V. Natarajan, Robust and data-efficient generalization of self-supervised machine learning for diagnostic imaging. Nat Biomed Eng 7, 756–779 (2023).

48. G. Arango-Argoty, E. Kipkogei, R. Stewart, G. J. Sun, A. Patra, I. Kagiampakis, E. Jacob, Pretrained transformers applied to clinical studies improve predictions of treatment efficacy and associated biomarkers. Nat. Commun. 16, 2101 (2025).

49. W. Shen, T. H. Nguyen, M. M. R. Li, Y. Huang, I. Moon, N. Nair, D. Marbach, M. Zitnik, Generalizable AI predicts immunotherapy outcomes across cancers and treatments, medRxiv (2025). 10.1101/2025.05.01.25326820.

50. T. J. Pugh, J. L. Bell, J. P. Bruce, G. J. Doherty, M. Galvin, M. F. Green, H. Hunter-Zinck, P. Kumari, M. L. Lenoue-Newton, M. M. Li, J. Lindsay, T. Mazor, A. Ovalle, S.-J. Sammut, N. Schultz, T. V. Yu, S. M. Sweeney, B. Bernard, AACR Project GENIE Consortium, Genomics and Analysis Working Group, AACR Project GENIE: 100,000 Cases and Beyond. Cancer Discov. 12, 2044–2057 (2022).

51. Cancer Genome Atlas Research Network, J. N. Weinstein, E. A. Collisson, G. B. Mills, K. R. M. Shaw, B. A. Ozenberger, K. Ellrott, I. Shmulevich, C. Sander, J. M. Stuart, The Cancer Genome Atlas Pan-Cancer analysis project. Nat. Genet. 45, 1113–1120 (2013).

52. A. Mayakonda, D.-C. Lin, Y. Assenov, C. Plass, H. P. Koeffler, Maftools: efficient and comprehensive analysis of somatic variants in cancer. Genome Res. 28, 1747–1756 (2018).

53. L. F. Spurr, M. Touat, A. M. Taylor, A. M. Dubuc, J. Shih, D. M. Meredith, W. V. Pisano, M. L. Meyerson, K. L. Ligon, A. D. Cherniack, Y. Y. Li, R. Beroukhim, Quantification of aneuploidy in targeted sequencing data using ASCETS. Bioinformatics 37, 2461–2463 (2021).

54. L. B. Alexandrov, J. Kim, N. J. Haradhvala, M. N. Huang, A. W. Tian Ng, Y. Wu, A. Boot, K. R. Covington, D. A. Gordenin, E. N. Bergstrom, S. M. A. Islam, N. Lopez-Bigas, L. J. Klimczak, J. R. McPherson, S. Morganella, R. Sabarinathan, D. A. Wheeler, V. Mustonen, PCAWG Mutational Signatures Working Group, G. Getz, S. G. Rozen, M. R. Stratton, PCAWG Consortium, The repertoire of mutational signatures in human cancer. Nature 578, 94–101 (2020).

55. A. Yaacov, G. Ben Cohen, J. Landau, T. Hope, I. Simon, S. Rosenberg, Cancer mutational signatures identification in clinical assays using neural embedding-based representations. Cell Rep Med 5, 101608 (2024).

56. G. Ciriello, M. L. Miller, B. A. Aksoy, Y. Senbabaoglu, N. Schultz, C. Sander, Emerging landscape of oncogenic signatures across human cancers. Nat. Genet. 45, 1127–1133 (2013).

57. T. Davoli, H. Uno, E. C. Wooten, S. J. Elledge, Tumor aneuploidy correlates with markers of immune evasion and with reduced response to immunotherapy. Science, doi: 10.1126/science.aaf8399 (2017).

58. E. L. Huttlin, R. J. Bruckner, J. Navarrete-Perea, J. R. Cannon, K. Baltier, F. Gebreab, M. P. Gygi, A. Thornock, G. Zarraga, S. Tam, J. Szpyt, B. M. Gassaway, A. Panov, H. Parzen, S. Fu, A. Golbazi, E. Maenpaa, K. Stricker, S. Guha Thakurta, T. Zhang, R. Rad, J. Pan, D. P. Nusinow, J. A. Paulo, D. K. Schweppe, L. P. Vaites, J. W. Harper, S. P. Gygi, Dual proteome-scale networks reveal cell-specific remodeling of the human interactome. Cell 184, 3022–3040.e28 (2021).

59. L. Csabai, D. Fazekas, T. Kadlecsik, M. Szalay-Bekő, B. Bohár, M. Madgwick, D. Módos, M. Ölbei, L. Gul, P. Sudhakar, J. Kubisch, O. J. Oyeyemi, O. Liska, E. Ari, B. Hotzi, V. A. Billes, E. Molnár, L. Földvári-Nagy, K. Csályi, A. Demeter, N. Pápai, M. Koltai, M. Varga, K. Lenti, I. J. Farkas, D. Türei, P. Csermely, T. Vellai, T. Korcsmáros, SignaLink3: a multi-layered resource to uncover tissue-specific signaling networks. Nucleic Acids Res 50, D701–D709 (2022).

60. P. Lo Surdo, M. Iannuccelli, S. Contino, L. Castagnoli, L. Licata, G. Cesareni, L. Perfetto, SIGNOR 3.0, the SIGnaling network open resource 3.0: 2022 update. Nucleic Acids Res 51, D631–D637 (2023).

61. H. Han, J.-W. Cho, S. Lee, A. Yun, H. Kim, D. Bae, S. Yang, C. Y. Kim, M. Lee, E. Kim, S. Lee, B. Kang, D. Jeong, Y. Kim, H.-N. Jeon, H. Jung, S. Nam, M. Chung, J.-H. Kim, I. Lee, TRRUST v2: an expanded reference database of human and mouse transcriptional regulatory interactions. Nucleic Acids Res 46, D380–D386 (2018).

62. P. V. Hornbeck, J. M. Kornhauser, S. Tkachev, B. Zhang, E. Skrzypek, B. Murray, V. Latham, M. Sullivan, PhosphoSitePlus: a comprehensive resource for investigating the structure and function of experimentally determined post-translational modifications in man and mouse. Nucleic Acids Res 40, D261–70 (2012).

63. Z. Li, S. Chen, J.-H. Jhong, Y. Pang, K.-Y. Huang, S. Li, T.-Y. Lee, UbiNet 2.0: a verified, classified, annotated and updated database of E3 ubiquitin ligase-substrate interactions. Database (Oxford) 2021 (2021).

64. X. Wang, Y. Li, M. He, X. Kong, P. Jiang, X. Liu, L. Diao, X. Zhang, H. Li, X. Ling, S. Xia, Z. Liu, Y. Liu, C.-P. Cui, Y. Wang, L. Tang, L. Zhang, F. He, D. Li, UbiBrowser 2.0: a comprehensive resource for proteome-wide known and predicted ubiquitin ligase/deubiquitinase-substrate interactions in eukaryotic species. Nucleic Acids Res 50, D719–D728 (2022).

65. J. S. Lee, A. Das, L. Jerby-Arnon, R. Arafeh, N. Auslander, M. Davidson, L. McGarry, D. James, A. Amzallag, S. G. Park, K. Cheng, W. Robinson, D. Atias, C. Stossel, E. Buzhor, G. Stein, J. J. Waterfall, P. S. Meltzer, T. Golan, S. Hannenhalli, E. Gottlieb, C. H. Benes, Y. Samuels, E. Shanks, E. Ruppin, Harnessing synthetic lethality to predict the response to cancer treatment. Nat Commun 9, 2546 (2018).

66. J. Wang, M. Wu, X. Huang, L. Wang, S. Zhang, H. Liu, J. Zheng, SynLethDB 2.0: a web-based knowledge graph database on synthetic lethality for novel anticancer drug discovery. Database (Oxford) 2022 (2022).

67. A. Kratz, M. Kim, M. R. Kelly, F. Zheng, C. A. Koczor, J. Li, K. Ono, Y. Qin, C. Churas, J. Chen, R. T. Pillich, J. Park, M. Modak, R. Collier, K. Licon, D. Pratt, R. W. Sobol, N. J. Krogan, T. Ideker, A multi-scale map of protein assemblies in the DNA damage response. Cell Syst 14, 447–463.e8 (2023).

68. D. Szklarczyk, R. Kirsch, M. Koutrouli, K. Nastou, F. Mehryary, R. Hachilif, A. L. Gable, T. Fang, N. T. Doncheva, S. Pyysalo, P. Bork, L. J. Jensen, C. von Mering, The STRING database in 2023: protein-protein association networks and functional enrichment analyses for any sequenced genome of interest. Nucleic Acids Res 51, D638–D646 (2023).

69. P. Veličković, G. Cucurull, A. Casanova, A. Romero, P. Liò, Y. Bengio, Graph Attention Networks, arXiv [stat.ML] (2017). http://arxiv.org/abs/1710.10903.

70. S. Brody, U. Alon, E. Yahav, How Attentive are Graph Attention Networks?, arXiv [cs.LG] (2021). http://arxiv.org/abs/2105.14491.

71. A. Vaswani, N. Shazeer, N. Parmar, J. Uszkoreit, L. Jones, A. N. Gomez, L. Kaiser, I. Polosukhin, Attention is all you need, arXiv [cs.CL] (2017). http://arxiv.org/abs/1706.03762.

72. J. Cheng, G. Novati, J. Pan, C. Bycroft, A. Žemgulytė, T. Applebaum, A. Pritzel, L. H. Wong, M. Zielinski, T. Sargeant, R. G. Schneider, A. W. Senior, J. Jumper, D. Hassabis, P. Kohli, Ž. Avsec, Accurate proteome-wide missense variant effect prediction with AlphaMissense. Science 381, eadg7492 (2023).

73. D. Zhang, W. Zhang, Y. Zhao, J. Zhang, B. He, C. Qin, J. Yao, DNAGPT: A generalized pre-trained tool for versatile DNA sequence analysis tasks, arXiv [q-bio.GN] (2023). http://arxiv.org/abs/2307.05628.

74. B. Vogelstein, E. R. Fearon, S. R. Hamilton, S. E. Kern, A. C. Preisinger, M. Leppert, Y. Nakamura, R. White, A. M. Smits, J. L. Bos, Genetic alterations during colorectal-tumor development. N. Engl. J. Med. 319, 525–532 (1988).

75. A. G. Robertson, J. Kim, H. Al-Ahmadie, J. Bellmunt, G. Guo, A. D. Cherniack, T. Hinoue, P. W. Laird, K. A. Hoadley, R. Akbani, M. A. A. Castro, E. A. Gibb, R. S. Kanchi, D. A. Gordenin, S. A. Shukla, F. Sanchez-Vega, D. E. Hansel, B. A. Czerniak, V. E. Reuter, X. Su, B. de Sa Carvalho, V. S. Chagas, K. L. Mungall, S. Sadeghi, C. S. Pedamallu, Y. Lu, L. J. Klimczak, J. Zhang, C. Choo, A. I. Ojesina, S. Bullman, K. M. Leraas, T. M. Lichtenberg, C. J. Wu, N. Schultz, G. Getz, M. Meyerson, G. B. Mills, D. J. McConkey, TCGA Research Network, J. N. Weinstein, D. J. Kwiatkowski, S. P. Lerner, Comprehensive Molecular Characterization of Muscle-Invasive Bladder Cancer. Cell 174, 1033 (2018).

76. Cancer Genome Atlas Network, Comprehensive molecular portraits of human breast tumours. Nature 490, 61–70 (2012).

77. Cancer Genome Atlas Research Network, Comprehensive genomic characterization of squamous cell lung cancers. Nature 489, 519–525 (2012).

78. M. L. Poeta, J. Manola, M. A. Goldwasser, A. Forastiere, N. Benoit, J. A. Califano, J. A. Ridge, J. Goodwin, D. Kenady, J. Saunders, W. Westra, D. Sidransky, W. M. Koch, TP53 mutations and survival in squamous-cell carcinoma of the head and neck. N. Engl. J. Med. 357, 2552–2561 (2007).

79. M. L. Gillison, W. M. Koch, R. B. Capone, M. Spafford, W. H. Westra, L. Wu, M. L. Zahurak, R. W. Daniel, M. Viglione, D. E. Symer, K. V. Shah, D. Sidransky, Evidence for a causal association between human papillomavirus and a subset of head and neck cancers. J. Natl. Cancer Inst. 92, 709–720 (2000).

80. Cancer Genome Atlas Research Network, Albert Einstein College of Medicine, Analytical Biological Services, Barretos Cancer Hospital, Baylor College of Medicine, Beckman Research Institute of City of Hope, Buck Institute for Research on Aging, Canada’s Michael Smith Genome Sciences Centre, Harvard Medical School, Helen F. Graham Cancer Center &Research Institute at Christiana Care Health Services, HudsonAlpha Institute for Biotechnology, ILSbio, LLC, Indiana University School of Medicine, Institute of Human Virology, Institute for Systems Biology, International Genomics Consortium, Leidos Biomedical, Massachusetts General Hospital, McDonnell Genome Institute at Washington University, Medical College of Wisconsin, Medical University of South Carolina, Memorial Sloan Kettering Cancer Center, Montefiore Medical Center, NantOmics, National Cancer Institute, National Hospital, Abuja, Nigeria, National Human Genome Research Institute, National Institute of Environmental Health Sciences, National Institute on Deafness &Other Communication Disorders, Ontario Tumour Bank, London Health Sciences Centre, Ontario Tumour Bank, Ontario Institute for Cancer Research, Ontario Tumour Bank, The Ottawa Hospital, Oregon Health &Science University, Samuel Oschin Comprehensive Cancer Institute, Cedars-Sinai Medical Center, SRA International, St Joseph’s Candler Health System, Eli &Edythe L. Broad Institute of Massachusetts Institute of Technology &Harvard University, Research Institute at Nationwide Children’s Hospital, Sidney Kimmel Comprehensive Cancer Center at Johns Hopkins University, University of Bergen, University of Texas MD Anderson Cancer Center, University of Abuja Teaching Hospital, University of Alabama at Birmingham, University of California, Irvine, University of California Santa Cruz, University of Kansas Medical Center, University of Lausanne, University of New Mexico Health Sciences Center, University of North Carolina at Chapel Hill, University of Oklahoma Health Sciences Center, University of Pittsburgh, University of São Paulo, Ribeir ão Preto Medical School, University of Southern California, University of Washington, University of Wisconsin School of Medicine &Public Health, Van Andel Research Institute, Washington University in St Louis, Integrated genomic and molecular characterization of cervical cancer. Nature 543, 378–384 (2017).

81. Cancer Genome Atlas Network, Comprehensive genomic characterization of head and neck squamous cell carcinomas. Nature 517, 576–582 (2015).

82. T. Sano, T. Oyama, K. Kashiwabara, T. Fukuda, T. Nakajima, Expression status of p16 protein is associated with human papillomavirus oncogenic potential in cervical and genital lesions. Am. J. Pathol. 153, 1741–1748 (1998).

83. D. Rischin, R. J. Young, R. Fisher, S. B. Fox, Q.-T. Le, L. J. Peters, B. Solomon, J. Choi, B. O’Sullivan, L. M. Kenny, G. A. McArthur, Prognostic significance of p16INK4A and human papillomavirus in patients with oropharyngeal cancer treated on TROG 02.02 phase III trial. J. Clin. Oncol. 28, 4142–4148 (2010).

84. T. Y. Seiwert, Z. Zuo, M. K. Keck, A. Khattri, C. S. Pedamallu, T. Stricker, C. Brown, T. J. Pugh, P. Stojanov, J. Cho, M. S. Lawrence, G. Getz, J. Brägelmann, R. DeBoer, R. R. Weichselbaum, A. Langerman, L. Portugal, E. Blair, K. Stenson, M. W. Lingen, E. E. W. Cohen, E. E. Vokes, K. P. White, P. S. Hammerman, Integrative and comparative genomic analysis of HPV-positive and HPV-negative head and neck squamous cell carcinomas. Clin. Cancer Res. 21, 632–641 (2015).

85. P. H. Kim, E. K. Cha, J. P. Sfakianos, G. Iyer, E. C. Zabor, S. N. Scott, I. Ostrovnaya, R. Ramirez, A. Sun, R. Shah, A. M. Yee, V. E. Reuter, D. F. Bajorin, J. E. Rosenberg, N. Schultz, M. F. Berger, H. A. Al-Ahmadie, D. B. Solit, B. H. Bochner, Genomic predictors of survival in patients with high-grade urothelial carcinoma of the bladder. Eur. Urol. 67, 198–201 (2015).

86. D. Miao, C. A. Margolis, N. I. Vokes, D. Liu, A. Taylor-Weiner, S. M. Wankowicz, D. Adeegbe, D. Keliher, B. Schilling, A. Tracy, M. Manos, N. G. Chau, G. J. Hanna, P. Polak, S. J. Rodig, S. Signoretti, L. M. Sholl, J. A. Engelman, G. Getz, P. A. Jänne, R. I. Haddad, T. K. Choueiri, D. A. Barbie, R. Haq, M. M. Awad, D. Schadendorf, F. S. Hodi, J. Bellmunt, K.-K. Wong, P. Hammerman, E. M. Van Allen, Genomic correlates of response to immune checkpoint blockade in microsatellite-stable solid tumors. Nat. Genet. 50, 1271–1281 (2018).

87. S. Mariathasan, S. J. Turley, D. Nickles, A. Castiglioni, K. Yuen, Y. Wang, E. E. Kadel III, H. Koeppen, J. L. Astarita, R. Cubas, S. Jhunjhunwala, R. Banchereau, Y. Yang, Y. Guan, C. Chalouni, J. Ziai, Y. Şenbabaoğlu, S. Santoro, D. Sheinson, J. Hung, J. M. Giltnane, A. A. Pierce, K. Mesh, S. Lianoglou, J. Riegler, R. A. D. Carano, P. Eriksson, M. Höglund, L. Somarriba, D. L. Halligan, M. S. van der Heijden, Y. Loriot, J. E. Rosenberg, L. Fong, I. Mellman, D. S. Chen, M. Green, C. Derleth, G. D. Fine, P. S. Hegde, R. Bourgon, T. Powles, TGFβ attenuates tumour response to PD-L1 blockade by contributing to exclusion of T cells. Nature 554, 544–548 (2018).

88. R. S. Vanguri, J. Luo, A. T. Aukerman, J. V. Egger, C. J. Fong, N. Horvat, A. Pagano, J. de A. B. Araujo-Filho, L. Geneslaw, H. Rizvi, R. Sosa, K. M. Boehm, S.-R. Yang, F. M. Bodd, K. Ventura, T. J. Hollmann, M. S. Ginsberg, J. Gao, MSK MIND Consortium, M. D. Hellmann, J. L. Sauter, S. P. Shah, Multimodal integration of radiology, pathology and genomics for prediction of response to PD-(L)1 blockade in patients with non-small cell lung cancer. Nat. Cancer 3, 1151–1164 (2022).

89. D. Liu, B. Schilling, D. Liu, A. Sucker, E. Livingstone, L. Jerby-Arnon, L. Zimmer, R. Gutzmer, I. Satzger, C. Loquai, S. Grabbe, N. Vokes, C. A. Margolis, J. Conway, M. X. He, H. Elmarakeby, F. Dietlein, D. Miao, A. Tracy, H. Gogas, S. M. Goldinger, J. Utikal, C. U. Blank, R. Rauschenberg, D. von Bubnoff, A. Krackhardt, B. Weide, S. Haferkamp, F. Kiecker, B. Izar, L. Garraway, A. Regev, K. Flaherty, A. Paschen, E. M. Van Allen, D. Schadendorf, Integrative molecular and clinical modeling of clinical outcomes to PD1 blockade in patients with metastatic melanoma. Nat. Med. 25, 1916–1927 (2019).

90. J. Jee, C. Fong, K. Pichotta, T. N. Tran, A. Luthra, M. Waters, C. Fu, M. Altoe, S.-Y. Liu, S. B. Maron, M. Ahmed, S. Kim, M. Pirun, W. K. Chatila, I. de Bruijn, A. Pasha, R. Kundra, B. Gross, B. Mastrogiacomo, T. J. Aprati, D. Liu, J. Gao, M. Capelletti, K. Pekala, L. Loudon, M. Perry, C. Bandlamudi, M. Donoghue, B. A. Satravada, A. Martin, R. Shen, Y. Chen, A. R. Brannon, J. Chang, L. Braunstein, A. Li, A. Safonov, A. Stonestrom, P. Sanchez-Vela, C. Wilhelm, M. Robson, H. Scher, M. Ladanyi, J. S. Reis-Filho, D. B. Solit, D. R. Jones, D. Gomez, H. Yu, D. Chakravarty, R. Yaeger, W. Abida, W. Park, E. M. O’Reilly, J. Garcia-Aguilar, N. Socci, F. Sanchez-Vega, J. Carrot-Zhang, P. D. Stetson, R. Levine, C. M. Rudin, M. F. Berger, S. P. Shah, D. Schrag, P. Razavi, K. L. Kehl, B. T. Li, G. J. Riely, N. Schultz, MSK Cancer Data Science Initiative Group, Automated real-world data integration improves cancer outcome prediction. Nature 636, 728–736 (2024).

91. H. B. Lengel, B. Mastrogiacomo, J. G. Connolly, K. S. Tan, Y. Liu, C. N. Fick, E. G. Dunne, D. He, M. B. Lankadasari, B. A. Satravada, Y. Sun, R. Kundra, C. Fong, S. Smith, G. J. Riely, C. M. Rudin, D. R. Gomez, D. B. Solit, M. F. Berger, B. T. Li, M. W. Mayo, I. Matei, D. C. Lyden, P. S. Adusumilli, N. Schultz, F. Sanchez-Vega, D. R. Jones, Genomic mapping of metastatic organotropism in lung adenocarcinoma. Cancer Cell 41, 970–985.e3 (2023).

92. T. Byrski, R. Dent, P. Blecharz, M. Foszczynska-Kloda, J. Gronwald, T. Huzarski, C. Cybulski, E. Marczyk, R. Chrzan, A. Eisen, J. Lubinski, S. A. Narod, Results of a phase II open-label, non-randomized trial of cisplatin chemotherapy in patients with BRCA1-positive metastatic breast cancer. Breast Cancer Res. 14, R110 (2012).

93. K. Shin, A. Lim, C. Zhao, D. Sahoo, Y. Pan, E. Spiekerkoetter, J. C. Liao, P. A. Beachy, Hedgehog signaling restrains bladder cancer progression by eliciting stromal production of urothelial differentiation factors. Cancer Cell 26, 521–533 (2014).

94. K. Shin, J. J. Lee, N. Guo, J. Kim, A. Lim, L. Qu, I. Mysorekar, P. Beachy, Hedgehog/Wnt feedback supports regenerative proliferation of epithelial stem cells in bladder. Nature 472, 110–114 (2011).

95. R. M. Samstein, C.-H. Lee, A. N. Shoushtari, M. D. Hellmann, R. Shen, Y. Y. Janjigian, D. A. Barron, A. Zehir, E. J. Jordan, A. Omuro, T. J. Kaley, S. M. Kendall, R. J. Motzer, A. A. Hakimi, M. H. Voss, P. Russo, J. Rosenberg, G. Iyer, B. H. Bochner, D. F. Bajorin, H. A. Al-Ahmadie, J. E. Chaft, C. M. Rudin, G. J. Riely, S. Baxi, A. L. Ho, R. J. Wong, D. G. Pfister, J. D. Wolchok, C. A. Barker, P. H. Gutin, C. W. Brennan, V. Tabar, I. K. Mellinghoff, L. M. DeAngelis, C. E. Ariyan, N. Lee, W. D. Tap, M. M. Gounder, S. P. D’Angelo, L. Saltz, Z. K. Stadler, H. I. Scher, J. Baselga, P. Razavi, C. A. Klebanoff, R. Yaeger, N. H. Segal, G. Y. Ku, R. P. DeMatteo, M. Ladanyi, N. A. Rizvi, M. F. Berger, N. Riaz, D. B. Solit, T. A. Chan, L. G. T. Morris, Tumor mutational load predicts survival after immunotherapy across multiple cancer types. Nat. Genet. 51, 202–206 (2019).

96. E. C. Pacheco-Pinedo, A. C. Durham, K. M. Stewart, A. M. Goss, M. M. Lu, F. J. Demayo, E. E. Morrisey, Wnt/β-catenin signaling accelerates mouse lung tumorigenesis by imposing an embryonic distal progenitor phenotype on lung epithelium. J. Clin. Invest. 121, 1935–1945 (2011).

97. H. Liu, K. Liu, Z. Dong, Targeting CDK12 for cancer therapy: Function, mechanism, and drug discovery. Cancer Res. 81, 18–26 (2021).

98. S. Yao, F. Meric-Bernstam, D. Hong, F. Janku, A. Naing, S. A. Piha-Paul, A. M. Tsimberidou, D. Karp, V. Subbiah, T. A. Yap, J. R. Ahnert, S. Pant, E. E. I. Dumbrava, C. Wathoo, E. Campbell, L. Yu, Y. Yamamura, S. Fu, Clinical characteristics and outcomes of phase I cancer patients with CCNE1 amplification: MD Anderson experiences. Sci. Rep. 12, 8701 (2022).

99. S. Qin, M. Yi, D. Jiao, A. Li, K. Wu, Distinct roles of VEGFA and ANGPT2 in lung adenocarcinoma and squamous cell carcinoma. J. Cancer 11, 153–167 (2020).

100. Q. Li, M. Jiang, S. Hong, J. Yang, X. Wu, J. Pang, Y. Chen, X. Zhao, X. Ding, Comprehensive genomic and clinical analyses identify APOBEC mutational signatures as a brain metastasis risk factor in lung adenocarcinoma patients. Transl. Oncol. 43, 101921 (2024).

101. A. Vasudevan, P. S. Baruah, J. C. Smith, Z. Wang, N. M. Sayles, P. Andrews, J. Kendall, J. Leu, N. K. Chunduri, D. Levy, M. Wigler, Z. Storchová, J. M. Sheltzer, Single-chromosomal gains can function as metastasis suppressors and promoters in colon cancer. Dev. Cell 52, 413–428.e6 (2020).

102. Cancer Genome Atlas Research Network, Comprehensive molecular profiling of lung adenocarcinoma. Nature 511, 543–550 (2014).

103. C. Zhou, Q. Li, C. Li, J. Yu, Y. Liu, G. Wang, K. Zhang, C. Ji, Q. Yan, L. He, H. Peng, J. Li, J. Wu, Z. Liu, P. Xie, C. Xiong, J. Pei, P. S. Yu, L. Sun, A comprehensive survey on pretrained Foundation Models: A history from BERT to ChatGPT, arXiv [cs.AI] (2023). http://arxiv.org/abs/2302.09419.

104. K. A. Hoadley, C. Yau, T. Hinoue, D. M. Wolf, A. J. Lazar, E. Drill, R. Shen, A. M. Taylor, A. D. Cherniack, V. Thorsson, R. Akbani, R. Bowlby, C. K. Wong, M. Wiznerowicz, F. Sanchez-Vega, A. G. Robertson, B. G. Schneider, M. S. Lawrence, H. Noushmehr, T. M. Malta, Cancer Genome Atlas Network, J. M. Stuart, C. C. Benz, P. W. Laird, Cell-of-Origin Patterns Dominate the Molecular Classification of 10,000 Tumors from 33 Types of Cancer. Cell 173, 291–304.e6 (2018).

105. M. Scheffner, B. A. Werness, J. M. Huibregtse, A. J. Levine, P. M. Howley, The E6 oncoprotein encoded by human papillomavirus types 16 and 18 promotes the degradation of p53. Cell 63, 1129–1136 (1990).

106. M. Yang, M. Wang, X. Li, Y. Xie, X. Xia, J. Tian, K. Zhang, A. Tang, Wnt signaling in cervical cancer? J. Cancer 9, 1277–1286 (2018).

107. T. Rampias, E. Boutati, E. Pectasides, C. Sasaki, P. Kountourakis, P. Weinberger, A. Psyrri, Activation of Wnt signaling pathway by human papillomavirus E6 and E7 oncogenes in HPV16-positive oropharyngeal squamous carcinoma cells. Mol. Cancer Res. 8, 433–443 (2010).

108. W. Choi, S. Porten, S. Kim, D. Willis, E. R. Plimack, J. Hoffman-Censits, B. Roth, T. Cheng, M. Tran, I.-L. Lee, J. Melquist, J. Bondaruk, T. Majewski, S. Zhang, S. Pretzsch, K. Baggerly, A. Siefker-Radtke, B. Czerniak, C. P. N. Dinney, D. J. McConkey, Identification of distinct basal and luminal subtypes of muscle-invasive bladder cancer with different sensitivities to frontline chemotherapy. Cancer Cell 25, 152–165 (2014).

109. D. J. McConkey, W. Choi, Y. Shen, I.-L. Lee, S. Porten, S. F. Matin, A. M. Kamat, P. Corn, R. E. Millikan, C. Dinney, B. Czerniak, A. O. Siefker-Radtke, A prognostic gene expression signature in the molecular classification of chemotherapy-naïve urothelial cancer is predictive of clinical outcomes from neoadjuvant chemotherapy: A phase 2 trial of dose-dense methotrexate, vinblastine, doxorubicin, and cisplatin with bevacizumab in urothelial cancer. Eur. Urol. 69, 855–862 (2016).

110. R. Seiler, H. A. D. Ashab, N. Erho, B. W. G. van Rhijn, B. Winters, J. Douglas, K. E. Van Kessel, E. E. Fransen van de Putte, M. Sommerlad, N. Q. Wang, V. Choeurng, E. A. Gibb, B. Palmer-Aronsten, L. L. Lam, C. Buerki, E. Davicioni, G. Sjödahl, J. Kardos, K. A. Hoadley, S. P. Lerner, D. J. McConkey, W. Choi, W. Y. Kim, B. Kiss, G. N. Thalmann, T. Todenhöfer, S. J. Crabb, S. North, E. C. Zwarthoff, J. L. Boormans, J. Wright, M. Dall’Era, M. S. van der Heijden, P. C. Black, Impact of Molecular Subtypes in Muscle-invasive Bladder Cancer on Predicting Response and Survival after Neoadjuvant Chemotherapy. Eur. Urol. 72, 544–554 (2017).

111. F. Zhong, Y. Lin, L. Zhao, C. Yang, Y. Ye, Z. Shen, Reshaping the tumour immune microenvironment in solid tumours via tumour cell and immune cell DNA methylation: from mechanisms to therapeutics. Br. J. Cancer 129, 24–37 (2023).

112. A. Chaudhri, G. Lizee, P. Hwu, K. Rai, Chromatin remodelers are regulators of the tumor immune microenvironment. Cancer Res. 84, 965–976 (2024).

113. N. Krishnamurthy, S. Kato, S. Lippman, R. Kurzrock, Chromatin remodeling (SWI/SNF) complexes, cancer, and response to immunotherapy. J. Immunother. Cancer 10, e004669 (2022).

114. M. Huang, F. Qian, Y. Hu, C. Ang, Z. Li, Z. Wen, Chromatin-remodelling factor BRG1 selectively activates a subset of interferon-alpha-inducible genes. Nat. Cell Biol. 4, 774–781 (2002).

115. A. H. Dudek, F. Pfaff, H. Bolte, C. Waguia Kontchou, M. Schwemmle, Partial inactivation of the chromatin remodelers SMARCA2 and SMARCA4 in virus-infected cells by caspase-mediated cleavage. J. Virol. 92 (2018).

116. M. Abou El Hassan, T. Yu, L. Song, R. Bremner, Polycomb repressive complex 2 confers BRG1 dependency on the CIITA locus. J. Immunol. 194, 5007–5013 (2015).

117. S. Papillon-Cavanagh, P. Doshi, R. Dobrin, J. Szustakowski, A. M. Walsh, STK11 and KEAP1 mutations as prognostic biomarkers in an observational real-world lung adenocarcinoma cohort. ESMO Open 5, e000706 (2020).

118. M. Knetki-Wróblewska, K. Wojas-Krawczyk, P. Krawczyk, M. Krzakowski, Emerging insights into STK11, KEAP1 and KRAS mutations: implications for immunotherapy in patients with advanced non-small cell lung cancer. Transl. Lung Cancer Res. 13, 3718–3730 (2024).

119. Cancer Genome Atlas Research Network. Electronic address, Cancer Genome Atlas Research Network, Integrated genomic characterization of pancreatic ductal adenocarcinoma. Cancer Cell 32, 185–203.e13 (2017).

120. Cancer Genome Atlas Research Network, The molecular taxonomy of primary prostate cancer. Cell 163, 1011–1025 (2015).

121. Cancer Genome Atlas Research Network. Electronic address, Cancer Genome Atlas Research Network, Comprehensive and integrated genomic characterization of adult soft tissue sarcomas. Cell 171, 950–965.e28 (2017).

122. J. Zhang, R. Bajari, D. Andric, F. Gerthoffert, A. Lepsa, H. Nahal-Bose, L. D. Stein, V. Ferretti, The international cancer genome consortium data portal. Nat. Biotechnol. 37, 367–369 (2019).

123. R. Kundra, H. Zhang, R. Sheridan, S. J. Sirintrapun, A. Wang, A. Ochoa, M. Wilson, B. Gross, Y. Sun, R. Madupuri, B. A. Satravada, D. Reales, E. Vakiani, H. A. Al-Ahmadie, A. Dogan, M. Arcila, A. Zehir, S. Maron, M. F. Berger, C. Viaplana, K. Janeway, M. Ducar, L. Sholl, S. Dogan, P. Bedard, L. F. Surrey, I. H. Sanchez, A. Syed, A. B. Rema, D. Chakravarty, S. Suehnholz, M. Nissan, G. V. Iyer, R. Murali, N. Bouvier, R. A. Soslow, D. Hyman, A. Younes, A. Intlekofer, J. J. Harding, R. D. Carvajal, P. J. Sabbatini, G. K. Abou-Alfa, L. Morris, Y. Y. Janjigian, M. M. Gallagher, T. A. Soumerai, I. K. Mellinghoff, A. A. Hakimi, M. Fury, J. T. Huse, A. Bagrodia, M. Hameed, S. Thomas, S. Gardos, E. Cerami, T. Mazor, P. Kumari, P. Raman, P. Shivdasani, S. MacFarland, S. Newman, A. Waanders, J. Gao, D. Solit, N. Schultz, OncoTree: A Cancer Classification System for Precision Oncology. JCO Clin Cancer Inform 5, 221–230 (2021).

124. S. Park, C.-Y. Ock, H. Kim, S. Pereira, S. Park, M. Ma, S. Choi, S. Kim, S. Shin, B. J. Aum, K. Paeng, D. Yoo, H. Cha, S. Park, K. J. Suh, H. A. Jung, S. H. Kim, Y. J. Kim, J.-M. Sun, J.-H. Chung, J. S. Ahn, M.-J. Ahn, J. S. Lee, K. Park, S. Y. Song, Y.-J. Bang, Y.-L. Choi, T. S. Mok, S.-H. Lee, Artificial Intelligence-Powered Spatial Analysis of Tumor-Infiltrating Lymphocytes as Complementary Biomarker for Immune Checkpoint Inhibition in Non-Small-Cell Lung Cancer. J. Clin. Oncol. 40, 1916–1928 (2022).

125. J. Kong, X. Zhao, A. Singhal, S. Park, R. Bachelder, J. Shen, H. Zhang, J. Moon, C. Ahn, C.-Y. Ock, H. Carter, T. Ideker, Prediction of immunotherapy response using mutations to cancer protein assemblies. Sci Adv 10, eado9746 (2024).

126. S. Park, E. Silva, A. Singhal, M. R. Kelly, K. Licon, I. Panagiotou, C. Fogg, S. Fong, J. J. Y. Lee, X. Zhao, R. Bachelder, B. A. Parker, K. T. Yeung, T. Ideker, A deep learning model of tumor cell architecture elucidates response and resistance to CDK4/6 inhibitors. Nat Cancer, doi: 10.1038/s43018-024-00740-1 (2024).

127. D. Pratt, J. Chen, D. Welker, R. Rivas, R. Pillich, V. Rynkov, K. Ono, C. Miello, L. Hicks, S. Szalma, A. Stojmirovic, R. Dobrin, M. Braxenthaler, J. Kuentzer, B. Demchak, T. Ideker, NDEx, the Network Data Exchange. Cell Syst 1, 302–305 (2015).

128. M. Fey, J. E. Lenssen, Fast graph representation learning with PyTorch Geometric, arXiv [cs.LG] (2019). http://arxiv.org/abs/1903.02428.

129. J. Devlin, M.-W. Chang, K. Lee, K. Toutanova, BERT: Pre-training of Deep Bidirectional Transformers for Language Understanding, arXiv [cs.CL] (2018). http://arxiv.org/abs/1810.04805.

130. A. Dosovitskiy, L. Beyer, A. Kolesnikov, D. Weissenborn, X. Zhai, T. Unterthiner, M. Dehghani, M. Minderer, G. Heigold, S. Gelly, J. Uszkoreit, N. Houlsby, An image is worth 16×16 words: Transformers for image recognition at scale, arXiv [cs.CV] (2020). http://arxiv.org/abs/2010.11929.

131. C. Ying, T. Cai, S. Luo, S. Zheng, G. Ke, D. He, Y. Shen, T.-Y. Liu, Do Transformers really perform bad for graph representation?, arXiv [cs.LG] (2021). http://arxiv.org/abs/2106.05234.

132. K. He, X. Zhang, S. Ren, J. Sun, “Deep residual learning for image recognition” in 2016 IEEE Conference on Computer Vision and Pattern Recognition (CVPR) (IEEE, 2016; 10.1109/cvpr.2016.90).

133. I. Loshchilov, F. Hutter, Decoupled Weight Decay Regularization, arXiv [cs.LG] (2017). http://arxiv.org/abs/1711.05101.

134. E. A. Eisenhauer, P. Therasse, J. Bogaerts, L. H. Schwartz, D. Sargent, R. Ford, J. Dancey, S. Arbuck, S. Gwyther, M. Mooney, L. Rubinstein, L. Shankar, L. Dodd, R. Kaplan, D. Lacombe, J. Verweij, New response evaluation criteria in solid tumours: revised RECIST guideline (version 1.1). Eur. J. Cancer 45, 228–247 (2009).

135. D. M. Gress, S. B. Edge, F. L. Greene, M. K. Washington, E. A. Asare, J. D. Brierley, D. R. Byrd, C. C. Compton, J. M. Jessup, D. P. Winchester, M. B. Amin, J. E. Gershenwald, “Principles of cancer staging” in AJCC Cancer Staging Manual (Springer International Publishing, Cham, 2017), pp. 3–30.

136. K. Clark, U. Khandelwal, O. Levy, C. D. Manning, What does BERT look at? An analysis of BERT’s attention, arXiv [cs.CL] (2019). http://arxiv.org/abs/1906.04341.

137. L. McInnes, J. Healy, J. Melville, UMAP: Uniform Manifold Approximation and Projection for Dimension Reduction, arXiv [stat.ML] (2018). http://arxiv.org/abs/1802.03426.

138. S. P. Suehnholz, M. H. Nissan, H. Zhang, R. Kundra, S. Nandakumar, C. Lu, S. Carrero, A. Dhaneshwar, N. Fernandez, B. W. Xu, M. E. Arcila, A. Zehir, A. Syed, A. R. Brannon, J. E. Rudolph, E. Paraiso, P. J. Sabbatini, R. L. Levine, A. Dogan, J. Gao, M. Ladanyi, A. Drilon, M. F. Berger, D. B. Solit, N. Schultz, D. Chakravarty, Quantifying the expanding landscape of clinical actionability for patients with cancer. Cancer Discov. 14, 49–65 (2024).

139. D. Chakravarty, J. Gao, S. M. Phillips, R. Kundra, H. Zhang, J. Wang, J. E. Rudolph, R. Yaeger, T. Soumerai, M. H. Nissan, M. T. Chang, S. Chandarlapaty, T. A. Traina, P. K. Paik, A. L. Ho, F. M. Hantash, A. Grupe, S. S. Baxi, M. K. Callahan, A. Snyder, P. Chi, D. Danila, M. Gounder, J. J. Harding, M. D. Hellmann, G. Iyer, Y. Janjigian, T. Kaley, D. A. Levine, M. Lowery, A. Omuro, M. A. Postow, D. Rathkopf, A. N. Shoushtari, N. Shukla, M. Voss, E. Paraiso, A. Zehir, M. F. Berger, B. S. Taylor, L. B. Saltz, G. J. Riely, M. Ladanyi, D. M. Hyman, J. Baselga, P. Sabbatini, D. B. Solit, N. Schultz, OncoKB: A precision oncology knowledge base. JCO Precis. Oncol. 2017 (2017).

140. F. Skoulidis, M. E. Goldberg, D. M. Greenawalt, M. D. Hellmann, M. M. Awad, J. F. Gainor, A. B. Schrock, R. J. Hartmaier, S. E. Trabucco, L. Gay, S. M. Ali, J. A. Elvin, G. Singal, J. S. Ross, D. Fabrizio, P. M. Szabo, H. Chang, A. Sasson, S. Srinivasan, S. Kirov, J. Szustakowski, P. Vitazka, R. Edwards, J. A. Bufill, N. Sharma, S.-H. I. Ou, N. Peled, D. R. Spigel, H. Rizvi, E. J. Aguilar, B. W. Carter, J. Erasmus, D. F. Halpenny, A. J. Plodkowski, N. M. Long, M. Nishino, W. L. Denning, A. Galan-Cobo, H. Hamdi, T. Hirz, P. Tong, J. Wang, J. Rodriguez-Canales, P. A. Villalobos, E. R. Parra, N. Kalhor, L. M. Sholl, J. L. Sauter, A. A. Jungbluth, M. Mino-Kenudson, R. Azimi, Y. Y. Elamin, J. Zhang, G. C. Leonardi, F. Jiang, K.-K. Wong, J. J. Lee, V. A. Papadimitrakopoulou, I. I. Wistuba, V. A. Miller, G. M. Frampton, J. D. Wolchok, A. T. Shaw, P. A. Jänne, P. J. Stephens, C. M. Rudin, W. J. Geese, L. A. Albacker, J. V. Heymach, STK11/LKB1 Mutations and PD-1 Inhibitor Resistance in KRAS-Mutant Lung Adenocarcinoma. Cancer Discov. 8, 822–835 (2018).

141. E. Cerami, J. Gao, U. Dogrusoz, B. E. Gross, S. O. Sumer, B. A. Aksoy, A. Jacobsen, C. J. Byrne, M. L. Heuer, E. Larsson, Y. Antipin, B. Reva, A. P. Goldberg, C. Sander, N. Schultz, The cBio cancer genomics portal: an open platform for exploring multidimensional cancer genomics data. Cancer Discov. 2, 401–404 (2012).

142. J. Gao, B. A. Aksoy, U. Dogrusoz, G. Dresdner, B. Gross, S. O. Sumer, Y. Sun, A. Jacobsen, R. Sinha, E. Larsson, E. Cerami, C. Sander, N. Schultz, Integrative analysis of complex cancer genomics and clinical profiles using the cBioPortal. Sci. Signal. 6, l1 (2013).

143. I. de Bruijn, R. Kundra, B. Mastrogiacomo, T. N. Tran, L. Sikina, T. Mazor, X. Li, A. Ochoa, G. Zhao, B. Lai, A. Abeshouse, D. Baiceanu, E. Ciftci, U. Dogrusoz, A. Dufilie, Z. Erkoc, E. Garcia Lara, Z. Fu, B. Gross, C. Haynes, A. Heath, D. Higgins, P. Jagannathan, K. Kalletla, P. Kumari, J. Lindsay, A. Lisman, B. Leenknegt, P. Lukasse, D. Madela, R. Madupuri, P. van Nierop, O. Plantalech, J. Quach, A. C. Resnick, S. Y. A. Rodenburg, B. A. Satravada, F. Schaeffer, R. Sheridan, J. Singh, R. Sirohi, S. O. Sumer, S. van Hagen, A. Wang, M. Wilson, H. Zhang, K. Zhu, N. Rusk, S. Brown, J. A. Lavery, K. S. Panageas, J. E. Rudolph, M. L. LeNoue-Newton, J. L. Warner, X. Guo, H. Hunter-Zinck, T. V. Yu, S. Pilai, C. Nichols, S. M. Gardos, J. Philip, AACR Project GENIE BPC Core Team, AACR Project GENIE Consortium, K. L. Kehl, G. J. Riely, D. Schrag, J. Lee, M. V. Fiandalo, S. M. Sweeney, T. J. Pugh, C. Sander, E. Cerami, J. Gao, N. Schultz, Analysis and visualization of longitudinal genomic and clinical data from the AACR Project GENIE Biopharma Collaborative in cBioPortal. Cancer Res. 83, 3861–3867 (2023).

